# Mechanism of action of HBV capsid assembly modulators predicted from binding to early assembly intermediates

**DOI:** 10.1101/2020.03.23.002527

**Authors:** Anna Pavlova, Leda Bassit, Bryan D. Cox, Maksym Korablyov, Chris Chipot, Kiran Verma, Olivia O. Russell, Raymond F. Schinazi, James C. Gumbart

## Abstract

Interfering with the self-assembly of virus nucleocapsids is a promising approach for the development of novel antiviral agents. Applied to hepatitis B virus (HBV), this approach has led to several classes of capsid assembly modulators (CAMs) that target the virus by either accelerating nucleocapsid assembly or misdirecting it into non-capsid-like particles. Here, we have assessed the structures of early nucleocapsid assembly intermediates, with and without bound CAMs, using molecular dynamics simulations. We find that distinct conformations of the intermediates are induced depending on whether the bound CAM accelerates or misdirects assembly; these structures are predictive of the final assembly. We also selected non-capsid-like structures from our simulations for virtual screening, resulting in the discovery of several compounds with moderate anti-viral activity and low toxicity. Cryo-electron microscopy and capsid melting experiments suggest that our compounds possess a novel mechanism for assembly modulation, potentially opening new avenues for HBV inhibition.

## Introduction

Hepatitis B virus (HBV) is the leading cause of liver complications, including cirrhosis, hepatocellular carcinoma and liver failure^1^. The chronic infection affects roughly 250 million people worldwide and approximately 800,000 individuals die every year from its complications^1^. It is characterized by a persistent presence of covalently closed circular DNA (cccDNA) in the infected hepatocytes, which is not eliminated by presently approved therapies^2–4^. A promising orthogonal approach for eliminating the infection is to target HBV nucleocapsid^5–13^. Capsid assembly modulators (CAMs) are small molecules that affect capsid assembly by interacting with the capsid proteins^5–10, 12, 13^. Different assembly effects, such as acceleration or misdirection, have been achieved by different CAMs^5–13^. It was also discovered that CAMs can both inhibit virus replication and interfere with cccDNA synthesis, suggesting that they could help eliminate the virus from hepatocytes more efficiently^5, 6^. However, a better understanding of HBV capsid assembly and the interference mechanisms of novel CAMs is needed for the rational development of more effective drugs.

HBV capsid protein, HBc, primary exists as a homodimer in solution^2^ and its N-terminal domain (Cp149) is sufficient for forming regular virus capsids^14^. The Cp149 dimer consists of two domains: the dimerization interface and the assembly interface (Figure 1A), with the latter interface forming inter-dimer contacts during capsid assembly^15, 16^. Previous studies proposed that the Cp149 dimers trigger capsid assembly by adopting an energetically unfavorable “assembly-active” conformation, which in turn leads to assembly nucleation (Figure 1C)^17–20^. It was also concluded that the assembly is nucleated by the formation of a hexamer, a triangular trimer of dimers, which is the rate-limiting step, and is followed by successive addition of dimers or small intermediates until the complete nucleocapsid is formed^17–19^.

**Figure 1.**
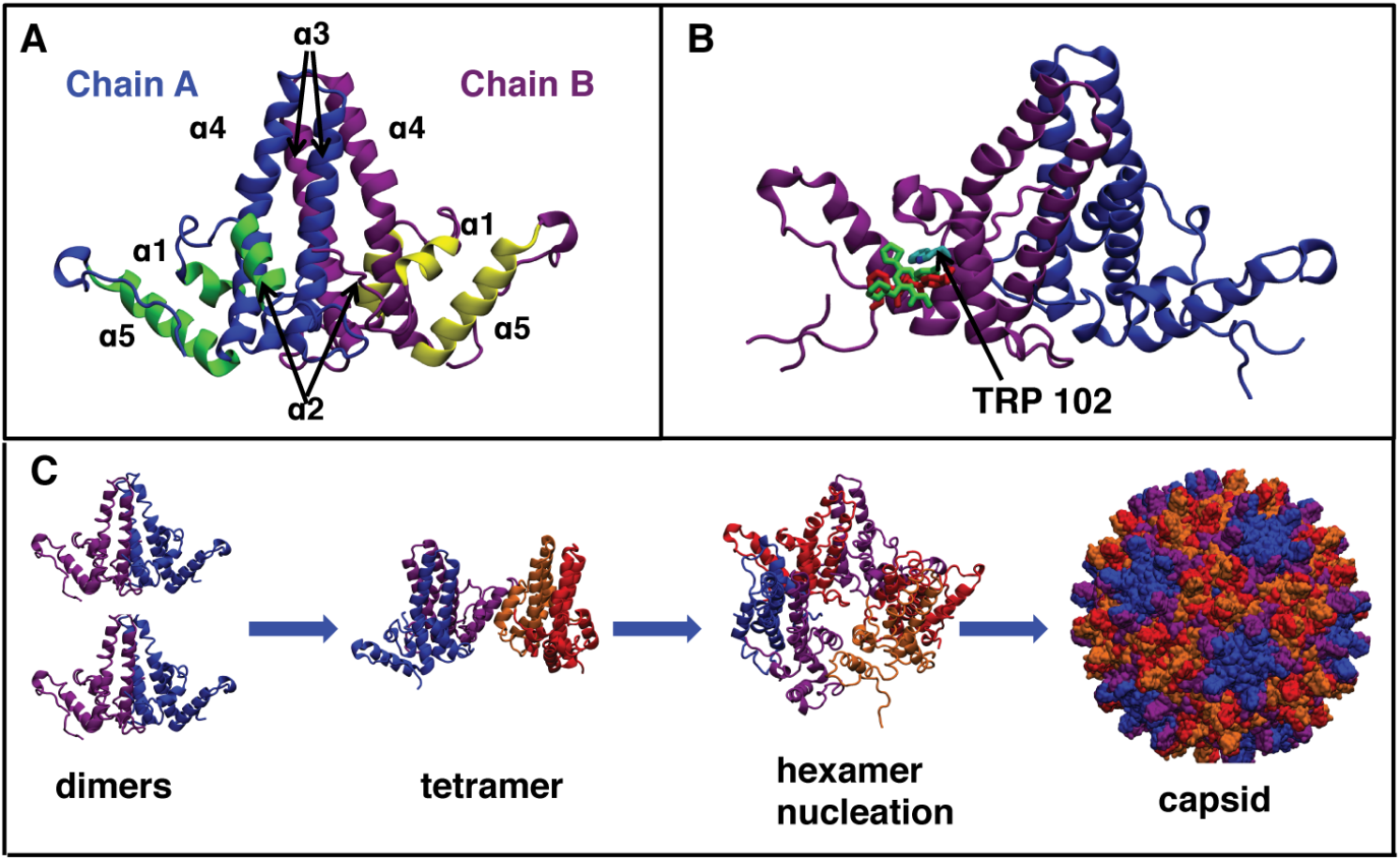
HBV capsid structure and assembly. (A) Structure of Cp149 dimer taken from the capsid structure (PDB code 3J2V). The names of helices *α*1-5 are indicated; yellow and green helices form the inter-dimer interface. (B) Overlap of GLS4 and AT130 binding sites after alignment of the bound protein structures (PDB codes 5E0I and 4G93, respectively.). Both compounds occupy a similar space in the HAP pocket and interact with Trp102. (C) HBV capsid assembly process based on experimental data and mathematical modeling^17, 27–29^.

Several factors, such as ions^17–19^, mutations^21–23^ and CAMs^5–10^ alter kinetics and/or thermodynamics of HBV capsid assembly, potentially preventing the formation of normal capsids. It has been suggested that the kinetic effects are caused by increased concentration of the “assembly-active” dimer conformation^18, 19^, while thermodynamic effects are caused by more favorable inter-dimer contact energies^6, 24, 25^. CAMs that alter HBV capsid assembly are divided into three structural and mechanistic classes (see Figure S1 for structures). Heteroaryldihydropyrimidines (HAPs) misdirect the capsid assembly into non-capsid structures^5, 6^, while phenylpropenamides (PPAs)^7, 8^ and sulfamoyl benzamides (SBAs)^9–11^ induce formations of capsids lacking the viral DNA. Although both PPAs and SBAs cause formation of empty capsids, it has been shown that some PPAs, e.g, AT130, also increase the assembly rate of Cp149^8^. In contrast, no changes were observed for Cp149 assembly with and without SBAs, suggesting that SBAs and PPAs alter the capsid assembly differently^9, 26^.

Crystal and cryogenic electron microscopy structures show that all CAMs bind in the same HAP pocket at the dimer-dimer interface (Figure 1B)^10, 24, 25, 30^. Slightly altered dimer–dimer orientations and several hydrophobic contacts between the CAM and protein residues in the pocket were observed, explaining the experimental thermodynamic effects^24, 25^. Nevertheless, further studies are needed to elucidate the enhanced assembly kinetics and the misdirection of assembly by HAP compounds. The known structures were obtained from CAMs binding to either pre-formed capsids or to a hexamer of the assembly-incompetent Y132A mutant^22^. However, it is possible that CAMs induce distinct structural changes in early assembly intermediates. Although these intermediates have been detected with mass spectroscopy^27–29^, little information is available about their structure, their dynamics and how these properties are altered upon CAM binding.

Both capsids and transient assembly intermediates can be studied with molecular dynamics (MD) simulations. It was previously shown that HAPs decrease the structural fluctuations of the Cp149 hexamer^31^ and flatten the hexameric units in the assembled capsid^32^. Additionally, prior MD simulations of Cp149 capsids showed that they are highly flexible, and that both CAMs and mutations can alter capsid dynamics^23, 33^. Here, we have assessed the structure and dynamics of Cp149 tetramers and hexamers in the presence of CAMs from the three known classes (HAPs, PPAs and SBAs) using MD simulations. Distinct structural changes in these intermediates were observed for the three classes of CAMs. Additionally, several structures from apo tetramer MD simulations were selected for docking; the top 58 candidate compounds were tested in HepAD38 cells. After two rounds of docking and testing we have discovered five structurally novel CAMs with low cytotoxicity and moderate activity against HBV (EC_50_≈10 *μ*M).

## Results

### Conformational changes in early assembly intermediates observed by MD

We performed MD simulations of a wild type (WT) Cp149 tetramer and a hexamer, as well as tetramers with either Y132A or V124W mutations, which inhibit or enhance capsid assembly, respectively^21, 22^. Although hepatitis B virus (HBV) nucleocapsids contain four quasi-equivalent tetramers and two quasi-equivalent hexamers, we have determined that the ABCD tetramer and the CDCDCD hexamer are the best starting points for tetramer and hexamer simulations (see Supporting Information for an explanation of nomenclature).

Significant differences in dimer-dimer orientation were observed for the hexamer in comparison to the tetramer (Table S3). These structural changes are well-described by changes in spike and base angles of the tetrameric unit (Figure 2D). The spike angle was calculated between the dimerization interfaces (*α*3 and *α*4 helices) of each dimer and describes the “bending” of the tetrameric unit. The base angle was calculated from the positions of the interface-forming *α*5 helices in each dimer and describes the “opening” and “closure” of the tetrameric unit. To illustrate the observed structural differences, the distributions of spike and base angles for each system were projected on a 2D scatter plot, and standard deviations ellipses (SDEs; Methods) were used to illustrate the spread of the distributions in our simulations^34^. The SDEs for the studied systems were quantitatively compared by calculating their fractional overlap areas (FOA) in Table S4. The values of FOA can range from 0% (no overlap) to 100% (perfect overlap). Values over 50% indicate notable structural similarity, while values close to 0% indicate significant structural differences.

**Figure 2.**
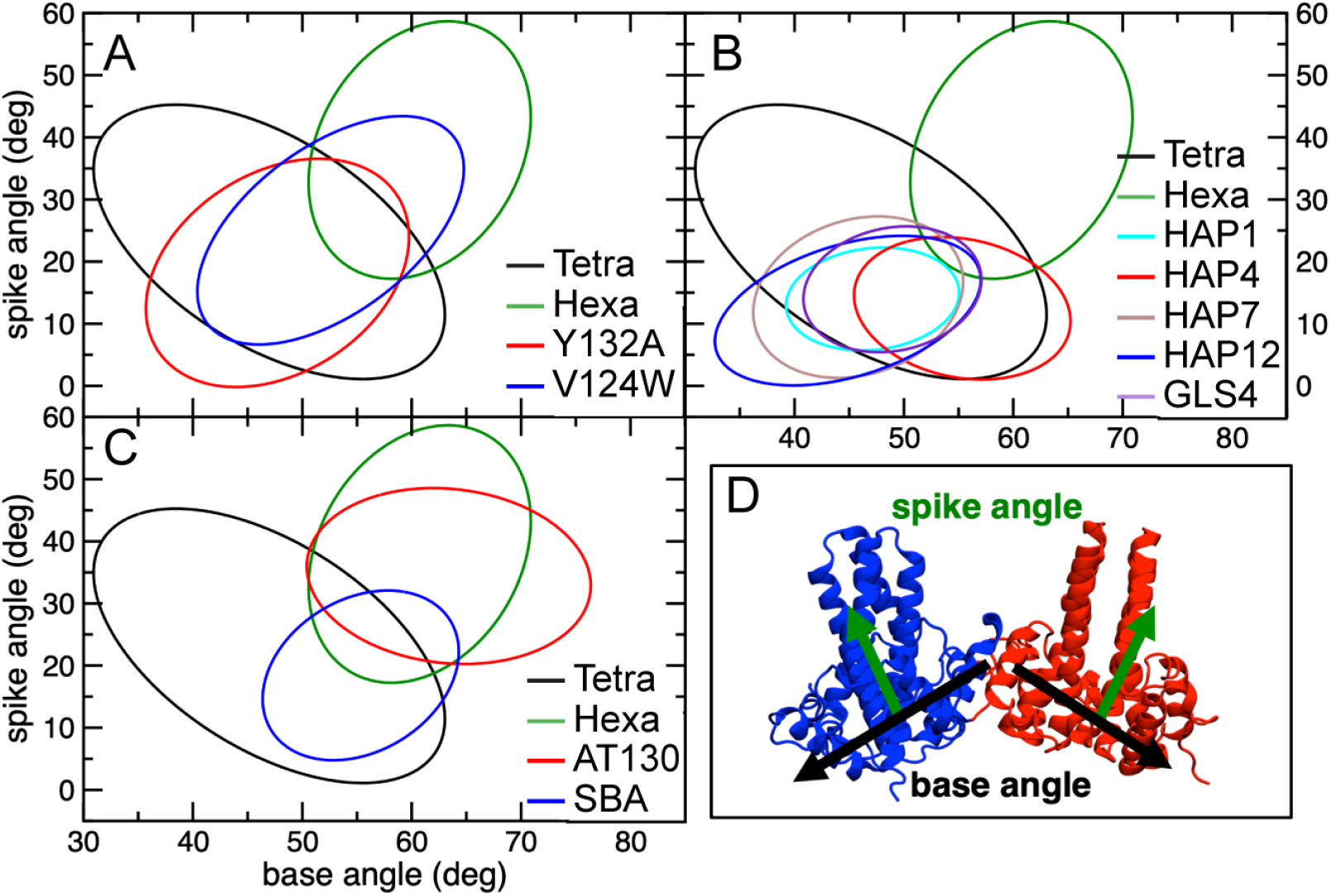
Comparison of standard deviation ellipses (SDEs) for spike and base angle distributions for selected simulations. SDEs are centered on the average values of a distribution. Their width and height are based on the standard deviation of the corresponding variables, while their rotation is based on the correlation between two variables. The ellipses shown here are scaled to envelop 90% of the sampled distributions. (A) SDEs of base and spike angles for the simulated apo-structures. (B) SDEs of base and spike angle for selected misdirectors (see Figure S9 for all misdirectors). The results for apo tetramer and hexamer are added for comparison. (C) SDEs of base and spike angles for SBA_R01, which does not alter the empty capsid assembly, and the accelerator AT130. The results for apo tetramer and hexamer are added for comparison. (D) The definitions of spike and base angles. See Methods and Supporting Information for details.

As shown in Figure 2A and Table S2, a wide range of base angles (31-63°) was observed for the tetramer. In comparison, the hexamer simulations displayed a more narrow range (49-72°), centered around 60°, as expected for a planar, symmetric hexamer. The hexamer also adopted larger spike angles than the tetramer (6-59° compared to 1-45°, respectively). The tetramer FOA with the hexamer is only 14%, showing that there are significant structural differences in terms of spike and base angles for the two systems. The FOA with the hexamer increases significantly to 48% for the assembly-enhancing V124W mutant tetramer (Figure 2A), which could decrease the energetic barrier for nucleus formation^21^. In contrast, the assembly-incompetent Y132A mutant tetramer^22^ showed similar FOA with the hexamer state as the WT tetramer (21%), suggesting that the assembly inhibition is not caused by altered inter-dimer orientation. We hypothesize that the “assembly active” conformation is a tetramer that adopts a more “hexamer-like” conformation, in agreement with the theory of allosteric assembly^18, 19, 35^. The frequency of such conformations is increased for the V124W mutant, explaining its acceleration of the capsid assembly kinetics. To ensure that our results are not dependent on the initial structures, we also simulated several systems using different starting states and arrived at the same conclusions (see Supporting Information).

### MD simulations show that HAPs and PPAs induce distinct conformational changes in early-assembly intermediates

In addition to simulations of the apo state, we have also simulated Cp149 tetramers with the following bound HAP compounds: BAY41-4109, HAP1, HAP4, HAP7, HAP12, and GLS4. GLS4 and HAP12 are the most potent HAP compounds, while HAP4 and HAP7 are some of the least potent ones (Table 2)^6, 26^. Furthermore, simulations of the PPA AT130, which accelerates capsid assembly^7, 8^, and compound SBA_R01, which does not alter empty capsid assembly^9, 10^, were performed. With the exception of HAP4, the remaining HAP compounds show very similar base and spike angle distributions with FOAs of 49-100% (Figure 2B). These distributions are narrow in the ranges of 32-58° and 0-25° for base and spike angles, respectively. Both angles are significantly smaller than in the case of the apo hexamer, while the spike angles are also smaller than for the apo tetramer (Tables S3). The HAP4-bound tetramer displayed spike angles similar to other HAPs, while the base angles were significantly higher (Table S3). No significant differences in structural distributions were observed between the less potent HAP7 and the most active HAPs, HAP12 and GLS4 (FOAs > 60%). In contrast, the structures observed in AT130 simulations are remarkably different from all HAPs (0% FOAs), with larger base (50-80°) and spike (20-50°) angles. These conformations are more “hexamer-like”, based on their overlap with the ones from apo hexamer simulations (76% FOA), explaining the assembly accelerating effects of AT130. Simulations of SBA_R01 bound to a tetramer showed a narrow distribution of base and spike angles within the ranges observed for apo WT tetramer (85% FOA), in agreement with the experimental results that showed this compound does not alter the assembly of empty capsids^9, 26^.

**Table 1.** List of simulated systems, BAY stands for BAY41-4109, tetra stands for tetramer and hexa stands for hexamer. The middle column indicates which pdb structure was used as a starting state, with the exception of our new compounds, which started from MD-generated conformations. The last column lists the total simulation time for each system; in aggregate, the simulation time is over 10 *μ*s.

| System | Structure | number of simulations<br>× time (ns) |
| --- | --- | --- |
| apo tetra | 3J2V | 800+1200+2×150 |
| apo hexa sym | 3J2V | 2×150 |
| apo hexa asym | 3J2V | 3×150 |
| tetra V124W | 3J2V | 2×150 |
| tetra Y132A | 3J2V | 2×150 |
| apo tetra | 5E0I | 2×150 |
| apo hexa | 5E0I | 2×150 |
| tetra Y132A | 5E0I | 2×150 |
| tetra with HAP1, HAP4<br>HAP7, HAP12, GLS4 or BAY | 5E0I | 2×150 |
| hexa with 3 GLS4 | 5E0I | 2×150 |
| hexa with 1 GLS4 | 5E0I | 2×150 |
| tetra with AT130 | 4G93 | 2×150 |
| hexa with 3 AT130 | 4G93 | 2×150 |
| hexa with 1 AT130 | 4G93 | 2×150 |
| tetra with SBA_R01 | 5T2P | 2×150 |
| tetra with GT-5, GT39<br>GT-40, GT-45, GT-46 or GT-47 | MD | 2×150 |

**Table 2.** Comparison of experimental binding free energies, relative to HAP12 (EC_50_ of 0.012 *μ*M^6^), to the ones calculated with FEP. The EC_50_s used for calculating experimental binding free energies according to Eq. 1 are also added. Compound structures are displayed in Figure S1, while Figure S12 shows the transformations used in FEP calculations.

| Name | $EC_{50}$<sup>6,26</sup><br>($\mu M$) | $\Delta\Delta G_{bind,HAP12}$<br>Exp (kcal/mol) | $\Delta\Delta G_{bind,HAP12}$<br>FEP (kcal/mol) |
| --- | --- | --- | --- |
| HAP1 | 0.13 | 1.5 | 1.6 |
| HAP4 | 1.9 | 3.1 | 2.6 |
| HAP7 | $\gg 10$ | $\gg 4$ | 1.4 |
| GLS4 | 0.001 | -1.5 | -1.1 |

We investigated if similar structural changes would be observed in hexamers with CAMs bound. In contrast to tetramers and apo-hexamers, these hexamers displayed a narrow range of base angles (55-65°) with either AT130 or GLS4 bound (Figure S11). However, the range of observed spike angles closely resembles those observed for the tetramers with the same bound compound (20-50° and 0-22° for AT130 and GLS4, respectively; see Table S3 for standard errors). Although a hexamer has three dimer interfaces, binding of only one CAM is sufficient to induce the observed conformational changes (Figure S10). Our results strongly suggest that the misdirecting and accelerating CAMs induce different structural changes in capsid proteins starting from early assembly intermediates, which could explain their distinct effects on capsid assembly. The reduced conformational sampling in CAM-bound hexamers suggests stabilization of the hexameric unit, in agreement with experimental data and previous simulations^6, 8, 31^.

Although standard MD simulations could elucidate the mechanistic differences between different classes of CAMs, the cause of different potencies within the same class was less clear. We hypothesized that different potencies could be attributed to different binding affinities for the core protein tetramer. Free energy perturbation (FEP) was used to investigate our hypothesis by calculating the differences in binding free energies (ΔΔ*G*_bind_) between several HAP compounds. These free-energy differences were compared to the ones calculated from experimental EC_50_s (Table 2)^6, 26^. We assumed that Boltzmann-weighted binding free energies are proportional to the experimental EC_50_s and, thus, used the equation:

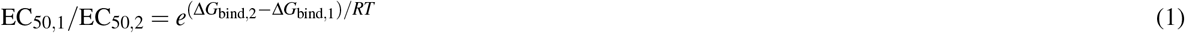

As shown in Table 2, there is very good agreement (within 0.6 kcal/mol) between calculated and experimentally estimated binding free energies for three out of four cases, indicating that these free energies are the largest factor responsible for the differences in the potency of HAP compounds. The only exception was HAP7 and HAP12, where experiments suggest a difference of ≫4 kcal/mol, while FEP predicted a difference of only 1.4 kcal/mol. It has been previously suggested that the protonation state is very important for CAMs targeting HBV and that binding of charged compounds should be less favorable due to the hydrophobic nature of the binding pocket^6^. Therefore, all our simulations employed the unprotonated, neutral states of each HAP compound. However, pK_a_ calculations using DFT (see Supporting Information) show that pK_a_s for HAP12 and HAP7 on the side chain nitrogen are 4.5 and 9.5 at 310 K, respectively. Therefore, only a fraction of HAP7 will be unprotonated at pH 7, explaining its low activity despite favorable binding free energy for the unprotonated state.

### Development of novel CAMs

We hypothesized that the knowledge of structural differences in early intermediates could be used for the development of novel CAMs and, therefore, extended two of our apo tetramer simulations for up to 1 *μ*s. Principal component analysis (PCA) was performed, with the top three components representing inter-dimer motions. Three structures from these simulations (Tetra1, Tetra2, and Tetra3) were selected for docking due to their large pocket volume and differences from capsid structures in PC space (Figure S13). Two databases were selected for the initial docking: the DivV set from NCI^36^ and ZINC0.9, which consists of all compounds with a Tanimoto similarity coefficient of 0.9 or lower in the ZINC database^37^. Both databases were docked to all three selected protein structures and the top 100 ranked compounds for each structure were considered for testing (see Methods for details on compound filtering). For the first round of experimental testing, we selected 29 compounds: 11 from the DivV set and 18 from the ZINC0.9 database (see Figures S15-S16 for structures and numbering).

The compounds were tested in HBVAD38 cells over seven days (see Methods for details). Table S6 shows the activity, toxicity and docking score for the tested compounds. Two compounds, GT-5 and GT-9, exhibited both activity against HBV and low toxicity (IC_50_ >50 *μ*M in all cell lines). They were probed for binding to capsid protein by measurement of tryptophan fluorescence, due to the presence of a tryptophan in the HAP pocket (Figure 1B). Only GT-5 showed a decrease in fluorescence in the presence of Cp149, suggesting direct binding to the protein (Figure 3). Size exclusion chromatography (SEC) also showed that GT-5 modestly promotes Cp149 assembly (Figure S20).

**Figure 3.**
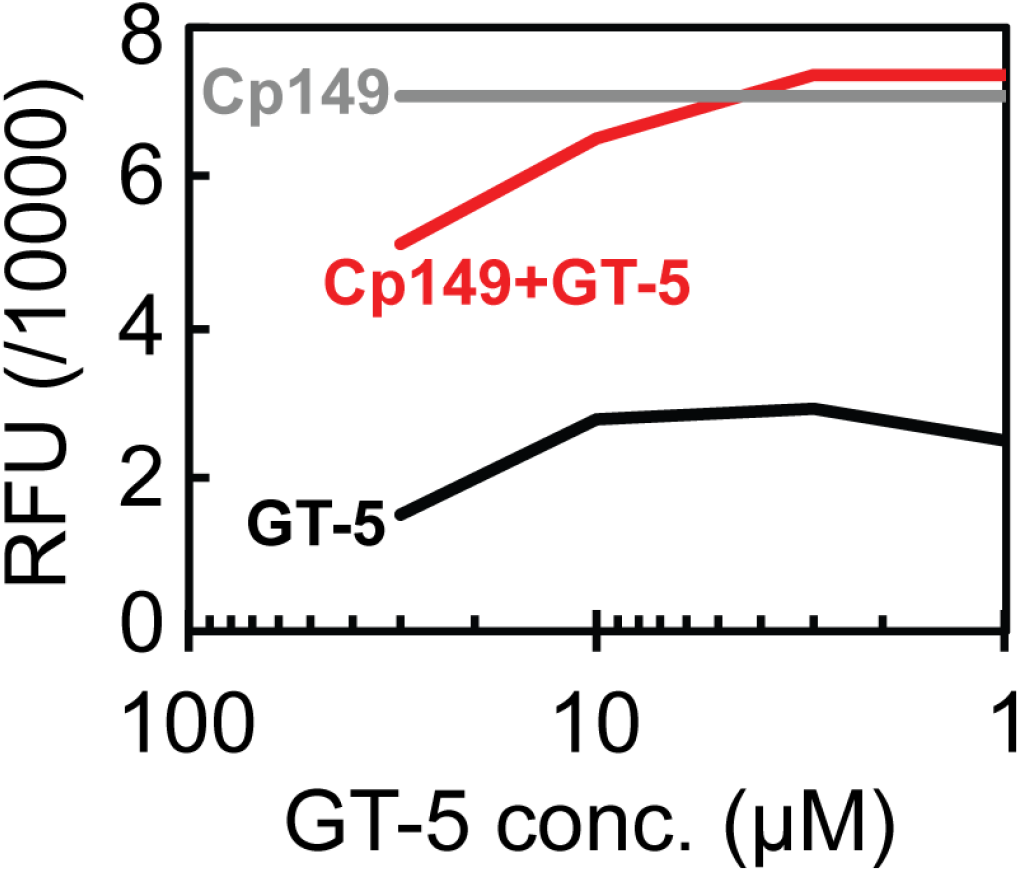
Measurement of HBV Cp149 tryptophan fluorescence in relative fluorescence units (RFU). The grey line shows data for HBV Cp149 dimer only in DMSO, the black line shows GT-5 only in buffer, and the red line shows GT-5 in the presence on Cp149 dimer. Measurements for GLS4 and GT-9 are shown in Figure S19.

Starting from GT-5, we performed a second round of docking using structurally similar compounds and protein structure Tetra2, for which GT-5 scored as one of the top compounds. NCI and Molport databases were searched for compounds with a similarity of at least 0.7 (Tanimoto coefficient)^36^. Additionally, compounds with a similar core as GT-5, but with different side-group substitutions, were investigated. In total, nearly 2000 different compounds were considered. Compounds with GlideXP docking scores lower than −8.0 kcal/mol were added to the list of potential new leads and filtered as described in Methods. Several compounds were removed from this list due to bad overlap with the docking pose of GT-5 based on visual inspection. In total 19 new compounds were selected for experimental testing (see Figures S17-S18 and Table S7). Seven of the tested compounds showed moderate inhibition of HBV DNA replication (∼50%) at 10 *μ*M concentration (Figure 4 and Table 3). Five of these compounds did not exhibit relevant toxicity at the effective concentrations, while two of them (GT-39 and GT-47) displayed low to moderate toxicity.

**Table 3.** Experimental and computational data for our most successful compounds. We display percentage inhibition (Inh.) of HBV DNA replication in HBVAD38 cells at 10 *μ*M compound concentration. Toxicity in four different cell types is also shown. Finally, changes in capsid melting temperature (ΔT_*m*_) are shown.

| name | HBV DNA | MTT cytotoxicity, $\text{IC}_{50}$ ($\mu\text{M}$) | | | | $\Delta T_m$ |
| --- | --- | --- | --- | --- | --- | --- |
| | Inh. (%)<br>at 10 $\mu\text{M}$ | PBM | CEM | Vero | HepG2 | $^{\circ}\text{C}$ |
| GT-5 | 24 | >100 | >100 | >100 | >100 | NA |
| GT-32 | 48 | 85 | 52 | 44 | 59 | $-0.9 \pm 0.0$ |
| GT-33 | 60 | >100 | >100 | 76 | 86 | NA |
| GT-39 | 57 | >100 | 16 | >100 | >100 | $-0.6 \pm 0.0$ |
| GT-40 | 55 | >100 | 77 | >100 | >100 | NA |
| GT-45 | 50 | >100 | >100 | 66 | >100 | $-0.5 \pm 0.2$ |
| GT-46 | 49 | >100 | 38 | 53 | 82 | $-2.9 \pm 0.0$ |
| GT-47 | 49 | >100 | >100 | >100 | 18 | $-1.2 \pm 0.4$ |

**Figure 4.**
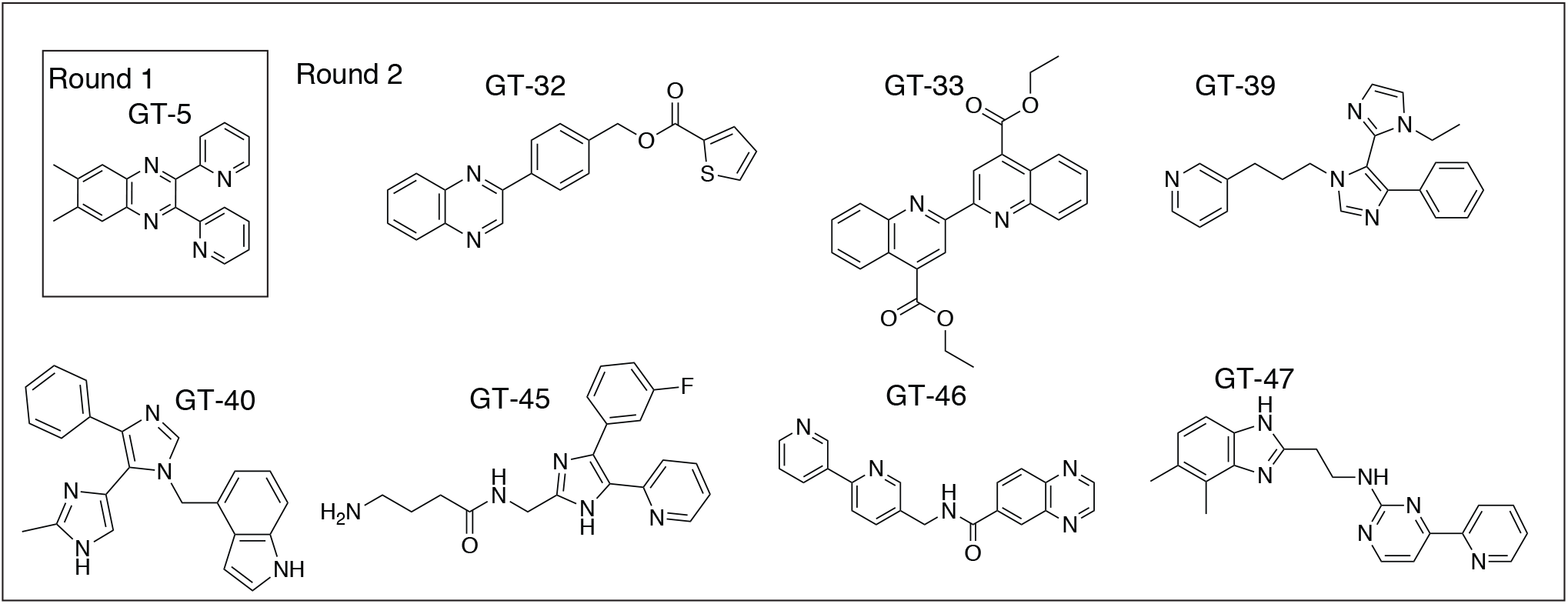
Structures of our compounds that showed moderate activity against HBV.

Thermal shift assays of Cp149 capsids in the presence of some of our compounds were performed to investigate if they altered capsid stability^30, 38^. Table 3 shows that GT-46 and GT-47 moderately decrease the capsid melting temperature (T_*m*_), suggesting they destabilize the capsid. The changes in T_*m*_ were not significant for the other tested compounds (T_*m*_ < 1.0 °C, Table 3). Cryo-electron microscopy (cryo-EM) imaging of preformed Cp149 capsids in the presence of GT-46 confirmed capsid destabilization by GT-46. Figure 5 shows that isolated Cp149 capsids had regular morphology (diameter~40 nm) and sparse cluster formation. In contrast, the addition of GT-46 resulted in fewer capsids with regular diameter and significant clustering of particles. In addition, the capsids appear misassembled or disintegrated (Figure 5).

**Figure 5.**
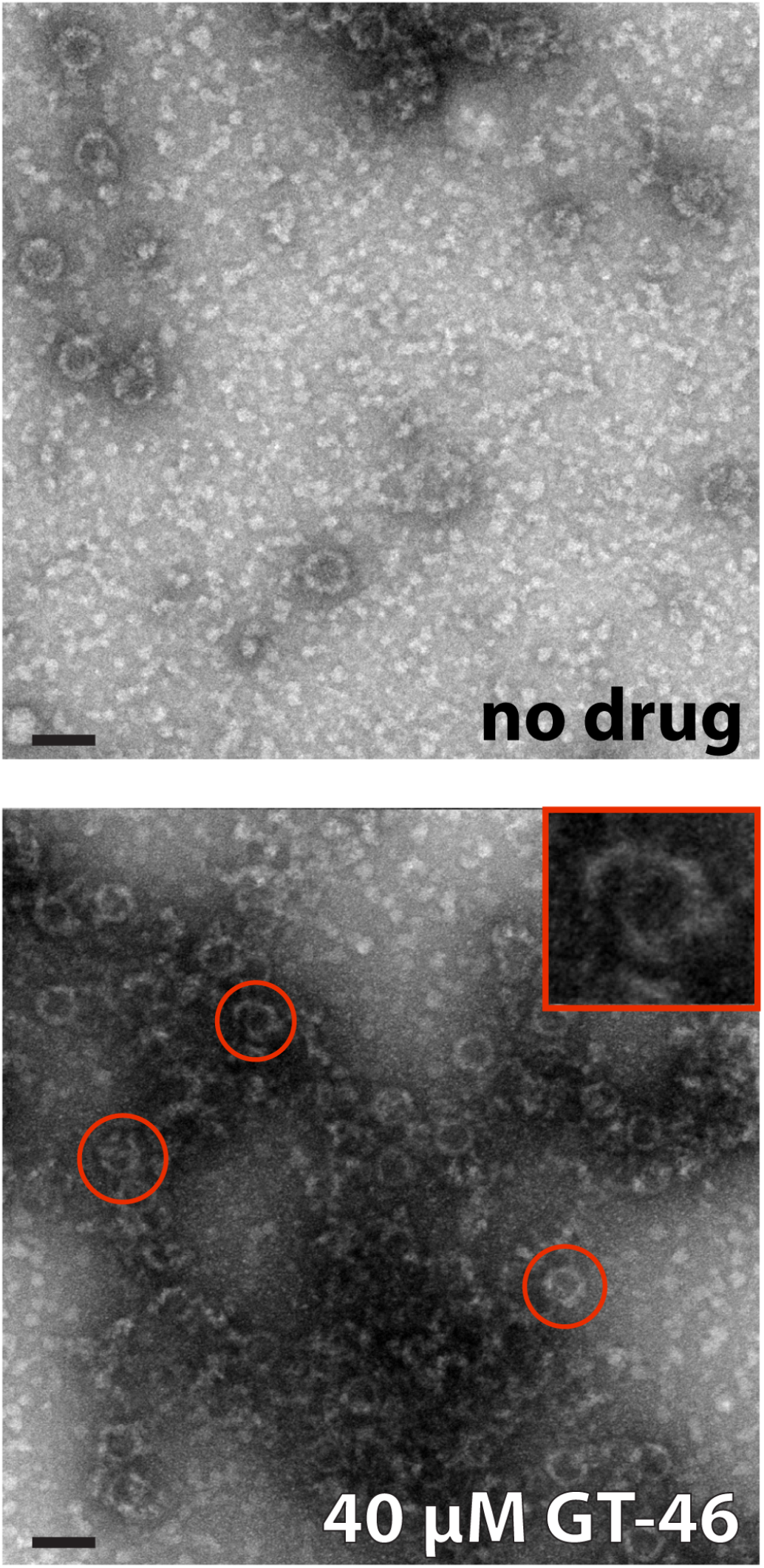
Selected images from EM of HBV Cp149 capsids with and without GT-46 (top and bottom images, respectively). A few examples of broken capsids in the presence of GT-46 are circled in the bottom image with one enlarged in the inset. The scale bar is 50 nm. Full images are presented in Figure S28.

We used MD simulations to investigate if and how our compounds induce distinct structural changes in early assembly intermediates. We simulated five of the most active compounds: GT-39, GT-40, GT-45, GT-46, and GT-47, as well as the initial lead GT-5. As shown in Figure 6, no clear trends in base and spike angles for these compounds are evident. GT-5 and GT-47 appear to accelerate the assembly due to larger spike angles (17-50° and 13-40°, respectively) in comparison to the apo tetramer (Table S3). In addition, their FOAs with the apo hexamer simulations are 71% (GT-5) and 59% (GT-47), which is a significant increase from the apo tetramer (14%). The simulations of GT-5 are in agreement with experimental data that showed it promotes assembly. In contrast, GT-39 and GT-40 simulations display base and spike angles similar to those of misdirecting compounds (Tables S2 and S3). Also, their FOAs with the HAP compounds are in the range 38-67%. Finally, GT-45 and GT-46 sample tetramer structures distinct from both accelerators and misdirectors with large base angles and relatively small spike angles (45-65° and 3-32°, respectively). Their FOAs with HAP compounds is ≤51% and even smaller with the apo hexamer ≤39%. Structurally, simulations of GT-45 and GT-46 resembled those of HAP4 (FOAs of 73% and 60%, respectively).

**Figure 6.**
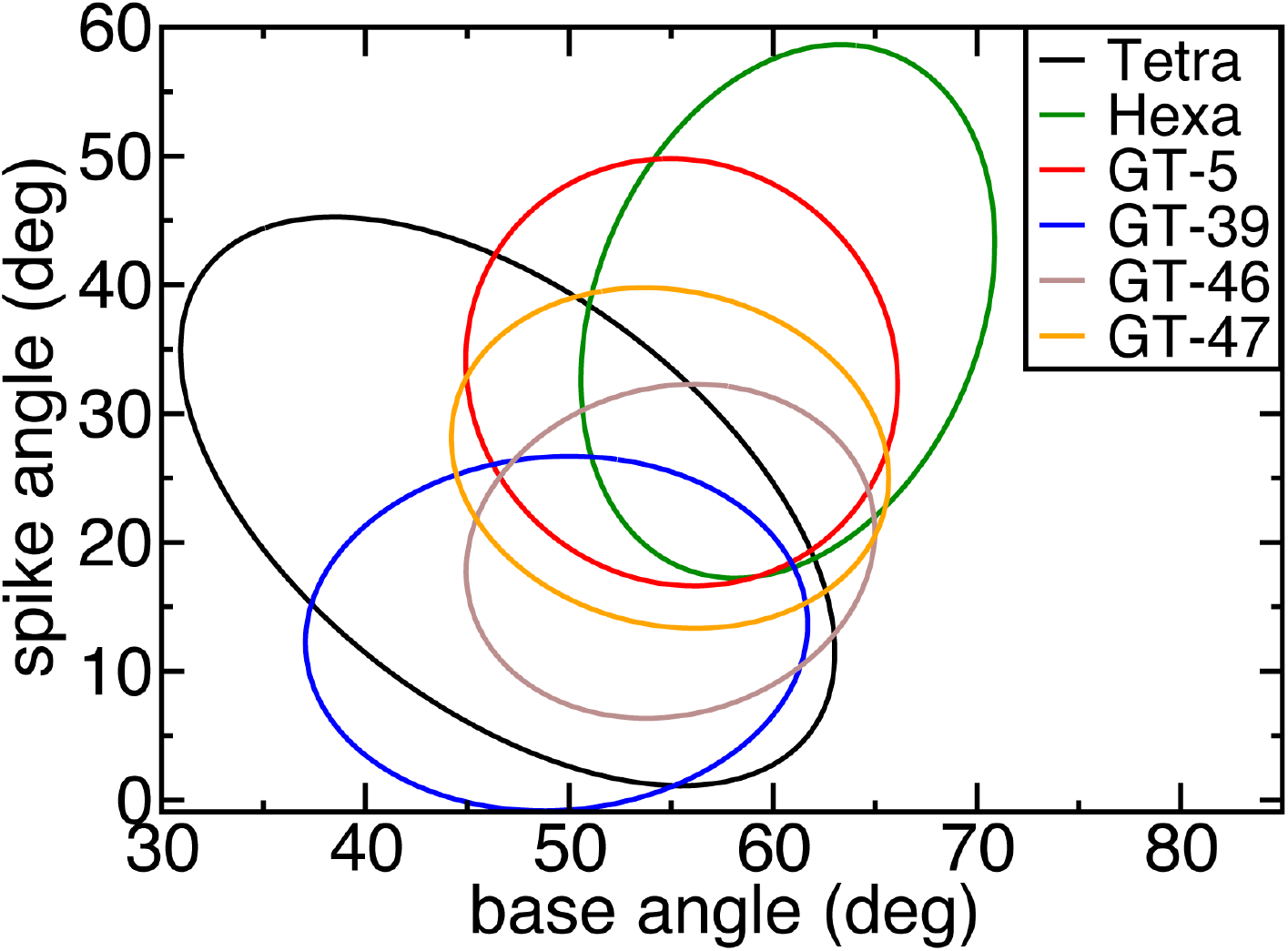
SDEs of base and spike angles for selected novel compounds. Apo tetramer and hexamer SDEs are added for comparison. The results for all novel compounds are shown in Figure S14.

## Discussion

Nucleocapsid assembly is an important event in the viral replication cycle^2, 12^. It is governed by weak association energies between core protein subunits and is highly sensitive to the assembly conditions^18, 35, 39^. It has been proposed that for HBV, the assembly is nucleated by an intermediate of three dimers, a triangular hexamer shown in Figure 1C, and that nucleus formation requires the adoption of a rare “assembly-active” conformation by a capsid protein dimer^17–19^. It should be noted that formation of a triangular hexamer from three “assembly-active” dimers encountering each other in solution is statistically unlikely. Furthermore, mass spectroscopy experiments are able to detect the formation of tetramers during capsid assembly, suggesting that they are also important assembly intermediates, and are likely hexamer precursors^27–29^. Based on what is known about HBV capsid assembly, there are three possibilities for the nucleation event: (1) addition of a third dimer to a tetramer, resulting in formation of an open hexamer, (2) closure of the open hexamer, or (3) a conformational change in the closed hexamer. Hexameric structures of the assembly-incompetent Y132A mutant have been crystallized in both open and closed states^20, 40^, suggesting that either hexamer closure or conformational change after this closure could be the bottleneck of the assembly process.

Our simulations revealed that tetramers and hexamers of the capsid protein sample different inter-dimer orientations and that the differences are well-described by base and spike angles between the dimers (Figures 2A and 2D). The tetramer exhibits greater structural flexibility than the hexamer, suggesting that only some of the tetramer conformations are able to incorporate another dimer to form an open hexamer, followed by hexamer closure. Based on our results, we propose that larger-than-average base and spike angles are required for transition from a tetramer to a hexamer. It has been shown that the dimer secondary structure does not change significantly during assembly^20, 41^; however, it is possible that minor structural changes govern assembly transitions. Because simulations of the hexamers starting from the capsid structure and the hexamer structure of the assembly incompetent Y132A mutant converge to similar distributions (see Figure S7), we propose that the nucleation event is a closure of the open hexamer, as opposed to a structural transition after hexamer closure.

CAMs can either accelerate capsid assembly or misdirect it into non-capsid structures^5–8^. HAP compounds lead to the formation of tubes and sheets instead of regular capsids^5, 6^, while AT130 causes the formation of regular capsids lacking viral DNA^7, 8^. Experiments have shown that both compound classes make dimer-dimer associations more energetically favorable and increase rates of nucleation and assembly^6, 42^. While crystal structures show stabilization of the dimer-dimer interfaces by CAMs through additional hydrophobic contacts^10, 25, 30^, they can not explain the observed kinetic effects nor how HAPs misdirect capsid assembly. Our simulations demonstrate that HAPs and AT130 introduce distinct changes to the structures of early assembly intermediates (Figures 2B and 2C). AT130 promotes structures with large base and spike angles, similar to the ones found for the apo hexamer. Such structures are highly curved and, therefore, are expected to favor formation of capsids (Figure 7).

**Figure 7.**
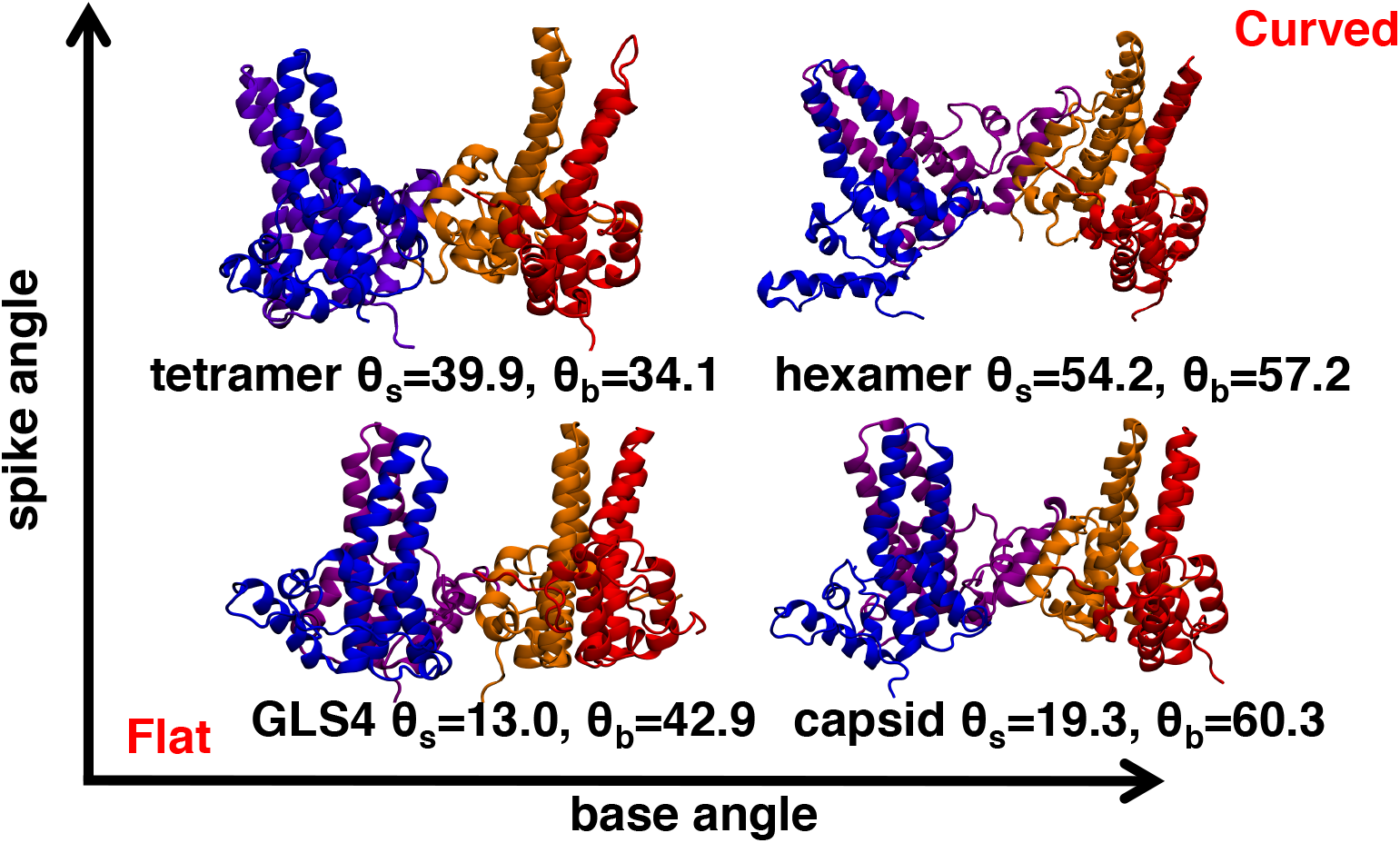
Selected snapshots of tetramers from our simulations illustrate the effect of spike and base angles on the curvature of the assembly intermediates.

In contrast, tetramers with bound HAP compounds sampled structures with smaller base and spike angles. Such structures are flat (Figure 7), explaining the formation of non-capsid structures in the presence of HAP compounds. Flattening of capsid-protein structures was also observed in the previous simulations of HBV capsids with bound HAP1^32^. Similar trends in curvature and spike angles are observed for tetramers and hexamers with the same CAM. However, hexamers with bound GLS4 or AT130 also sampled a narrower range of base angles, suggesting stabilization of the hexameric nucleus for both compounds. The observation that AT130 and GLS4 stabilize the hexameric nuclei in two distinct conformations supports our conclusion that the nucleation is related to hexamer closure and not to subsequent structural transitions. Our results could be confirmed by determining the collision cross-section (Ω) of intermediates in the presence of misdirecting and accelerating CAMs, as was previously done by Uetretcht et al. for intermediates during unmodified assembly^27^. Based on our data, we propose distinct changes in Ω for assembly intermediates in the presence of accelerating and misdirecting CAMs.

Finally, relative binding free energies, calculated using free energy perturbation (FEP), are highly correlated with experimental EC_50_s (Table 2), indicating that HAP potency is related to binding affinity and not to differences in induced structures. Based on our results, we propose that standard MD simulations and structural analysis can be used for predicting the mode of action of novel CAMS, while FEP can be used for predicting changes in CAM potency due to smaller structural modifications. Our observations regarding structural differences between apo and drug-bound tetramers as well as apo-hexamers were used to select tetramer structures for MD for docking. Multiple active compounds were identified by docking and subsequent experimental testing. The identified compounds appear to have a mechanism distinct from previous known CAMs against HBV. Our initial lead GT-5 is an assembly accelerator according to size-exclusion chromatography and MD simulations. In contrast, GT-45 and GT-46 have a novel mechanism of action, in which they destabilize the HBV nucleocapsid. In agreement with experimental data, MD simulations showed that tetramers with GT-45 and GT-46 bound adopt structures distinct from the ones observed in presence of known misdirectors and accelerators (Figure 6, Table S4). Our results illustrate that CAMs with a wide range of effects can be developed by a combined MD/docking approach, and that MD simulations of assembly intermediates with bound CAMs can predict their mechanism of action.

## Conclusions

Our results show that tetrameric and hexameric nucleocapsid assembly intermediates of HBV adopt distinct tertiary structures, which limits the rate of the capsid assembly. We propose that assembly nucleation is initiated by capsid-protein tetramers adopting a “hexamer-like” conformation, characterized by larger base and spike angles. Certain mutations as well as binding of assembly-accelerating CAMs increase the frequency of such tetramer conformations. In contrast, structure-misdirecting CAMs induce the formation of flat assembly intermediates with low spike angles, which can explain their effects on assembly. Our observations were used to select structures from MD simulations for docking, resulting in the development of several structurally novel CAMs. Experimental results suggest that our compounds may operate by a different mechanism than previously known CAMs. Our work demonstrates the importance of studying early capsid assembly intermediates, in particular their inter-subunit motions, in order to better understand the assembly of nucleocapsids and how to interfere with it.

## Methods

### Molecular Dynamics

The following structures were used in our simulations: 3J2V^43^, 4G93^25^, 5E0I^30^, and 5T2P^10^. 3J2V is the latest structure of the wild type (WT) HBV capsid, 4G93 is the structure of AT130 bound to pre-formed HBV capsids, in which several native cysteines were mutated to alanines, while the last two structures are of the hexameric Y132A mutant with bound drugs. 5E0I was crystallized with bound NVR-010-001-E2, which differs from GLS4 by a missing methyl group (Figure S1), whereas 5T2P was crystallized with bound SBA_R01. Structure preparation and MD protocols are described in Supporting Information. Table 1 displays which structure was used for each simulated system. In total 26 simulation systems were constructed for ~10 *μ*s of MD simulations.

NAMD2.12^44^ was used for the simulations of apo structures and the tetramers with bound existing CAMs, while AMBER16^45^ was used for simulations of our novel compounds and second runs of hexamer and tetramer simulations. The CHARMM36 force field was employed for all systems^46^. The energies of all systems were minimized before equilibration. For apo systems, the energy of all atoms was minimized at once, while for the systems with bound compounds a two-step minimization was used. In the first energy minimization step only water and ions were unrestrained, followed by an unrestrained energy minimization for all the atoms. Previously, we found that a two-step minimization can increase the compound stability in the binding pocket^47^. After minimization, a two-step equilibration was preformed for all systems. First, water and ions were equilibrated for 0.5 ns while restraining the protein and the CAM. In the second, 1-ns-long equilibration step, the restraints were removed from the CAM and protein side chains. Harmonic force constants of 2 kcal·mol^−1^ Å^−2^ were used for restraints in all cases. See Table 1 for the length and number of production runs for each system.

### Free energy perturbation (FEP) calculations

The relative binding free energies of a series of four substrates, namely HAP1, HAP4, HAP7 and GLS4, to Cp149 was determined with respect to HAP12 using the free energy perturbation (FEP) method^48, 49^. Towards this end, point mutation of the substrates was carried out in bulk water (unbound state) and at the binding site (bound state). Considering the nature of the point mutations, the reaction path was stratified into 50 stages of equal widths. Each alchemical transformation was run for 15 ns in the unbound state, and for 15 ns in the bound state, except for the transformation of HAP12 into HAP7, for which sampling was increased to 40 ns in bulk water, and 80 ns in the protein (see Table S5). The dual-topology paradigm was utilized, whereby a common scaffold is sought, and the chemical moieties characteristic of the initial and the final states of the transformation coexist but do not interact^50^. See Supporting Information for additional details.

### Analysis of simulations

The first 10 ns of each production run was discarded prior to analysis, after which the trajectory frames were analyzed with a frequency of 0.5 ns. The following definitions were used for base and spike angles: the base angle was calculated based on the positions of the *α*5 helices, while the spike angle was calculated based on the positions of *α*3 and *α*4 helices (see Supporting Information). Because the top parts of the helices *α*3 and *α*4 were very flexible, only the bottom parts of these helices were used for spike angle calculations, as described in the Supporting Information. Geometric centers of backbone atoms were used for all base and spike calculations. For each system, the data from 2 × 150 ns simulations was combined and projected on a 2D scatter plot. Standard deviation ellipses (SDEs) were drawn for each system to enable easier comparison of sampled structures. In a 2D plot SDEs are centered at the average values of the two variables, while the relative height and width are determined by the standard deviations of these variables^34^. The rotation of the SDE is calculated from variable correlation, and the total ellipse size is scaled to encompass a specific percentage of the provided distribution^34^, which corresponds to the confidence level of the ellipse. We choose to plot ellipses corresponding to 90% confidence level^34^. The distributions of base and spike angles are shown as scatter plots in Figures S24-S27. Additionally, comparison of SDEs between the two simulations of each system are shown in Figures S21-S23.

### Docking

All docking was done with Glide from Schrödinger suite with the default settings^51, 52^. We used Glide high-throughput virtual screening (HTVS) for the initial screening on the ZINC0.9 library^37, 51^. The 20000 highest scoring compounds were selected for re-docking with Glide single precision (SP), and the top 5000 of those compounds were re-docked with Glide extra precision (XP)^53^. For the DivV library, we first used Glide SP to dock all the ligands and selected the top 1000 ligands for re-docking with Glide XP. During optimization of our hit compound, we only needed to dock around 2000 compounds and therefore, only used Glide XP^51, 53^. When selecting top scoring molecules, compounds with expected low solubility, high reactivity, or toxicity, were discarded. Molecules that contained PAINS groups^54^ or were not readily available to order were excluded as well. Finally, we aimed for structural variance and to select compounds from different databases for testing.

### Inhibition assays

HBVAD38 cells were seeded at 50,000 cells/well in collagen-coated 96-well plates. Test compounds were added to HBVAD38 cells to a final concentration of 10 *μ*M. The experiment lasted 7 days. On day 7, total DNA was purified from supernatant using commercially available kit (DNeasy 96 Blood & Tissue kit, Qiagen). The HBV DNA was amplified in a real-time PCR assay using LightCycler 480 (Roche) as previously described^55^. All samples were tested in duplicate. Analysis: The concentration of compound that inhibited HBVDNA replication by 50% (EC_50_) was determined by linear regression.

### Cytotoxicity assays

Primary blood mononuclear (PBM), T lymphoblast CEM-CCRF (herein referred to as CEM cells), African green monkey kidney (Vero) or human liver (HepG2) cells were performed via MTT assay using the CellTiter 96 Non-Radioactive Cell Proliferation (Promega) kit as previously described^55^. Cytotoxicity was expressed as the concentration of test compounds that inhibited cell proliferation by 50 % (IC_50_) and calculated using the Chou and Talalay method^56^.

### Measurement of TRP fluorescence

The drugs were titrated at 3-fold dilutions in carbonate dimer buffer (pH=9.5, no NaCl). Then, HBV Cp149 dimer was added to a final concentration of 2 *μ*M and a volume of 100 *μ*L Intrinsic TRP fluorescence was recorded on a Biotek Cytation 3.0 with ex/em 285/350nm.

### Monitoring of capsid assembly

We incubated 20 *μ*M HBV Cp149 dimer with 40 *μ*M drug for 20 min, vol = 0.5 mL. 0.5 mL of SEC buffer (50mM Tris, 200 mM NaCl) was added and 0.7 mL of sample was immediately loaded onto Sephadex 16/60 gel filtration column and eluted at 1mL/min. 1 data set = 3 hours.

### Thermal shift assays

Assays were performed as previously described by Klumpp et al.^30^. Concentrations of 4 *μ*M HBV core protein monomer and 25 *μ*M ligand were used, as well as 1% DMSO. Melting temperatures were measured after 1 h incubation. The melting temperature of capsid without added compounds was 69.7 ± 0.5°C. All assays were performed twice, and the standard deviation from the two resulting values was used as an error estimate.

### EM imaging

HBV Cp149 protein was expressed in BL21 *E*.coli and isolated using established chromatographic methods^11^. Capsid particle formation was induced by decreasing pH and addition of NaCl overnight. We added 40 *μ*M of GT-46 to a sample of 5 *μ*M preformed capsids. This concentration was chosen as it is log(0.5) units greater than the antiviral EC50 (∼10 *μ*M). HBV Cp149 capsid assemblies were fixed onto a charged carbon grid and stained by uranyl acetate contrast agent for 15 min. EM images were collected using a JEOL JEM-1400 electron microscope operating at 120 kV at 25,000–35,000× magnification (Emory University Robert P. Apkarian electron microscopy core facility).

## Acknowledgements

This work was supported in part by NIH Grant 1-R01-AI-132833, and 5P30-AI-50409 (CFAR). Computational resources were provided via the Extreme Science and Engineering Discovery Environment (XSEDE; allocation TG-MCB130173), which is supported by NSF grant number ACI-1548562. Additional resources were provided by the Partnership for an Advanced Computing Environment (PACE) at the Georgia Institute of Technology.

