## Supplemental Information for "Mechanism of action of HBV capsid assembly modulators predicted from binding to early assembly intermediates"

### Supporting informations for "Mechanism of action of HBV capsid assembly modulators can be predicted from binding to early assembly intermediates"

**Anna Pavlova[1], Leda Bassit[2], Bryan D. Cox[2], Maksym Korablyov[3], Chris Chipot[4,5], Kiran Verma[2], Olivia O. Russell[2], Raymond F. Schinazi[2], and James C. Gumbart[1,*]**

[1]School of Physics and School of Chemistry & Biochemistry, Georgia Institute of Technology, Atlanta, GA, USA
[2]Center for AIDS Research, Laboratory of Biochemical Pharmacology, Department of Pediatrics, Emory University School of Medicine, Atlanta, Georgia, USA
[3]MIT Media Lab, Massachusetts Institute of Technology, Boston, MA, USA
[4]Department of Physics, University of Illinois at Urbana-Champaign, Urbana, IL, USA
[5]Laboratoire international associè CNRS-UIUC. UMR 7019 Université de Lorraine. B.P. 70239. 54506 Vandœuvre-lès-Nancy, France
[*]

#### Computational Details

##### Structure Preparation

For structures containing mutations, the mutations back to the WT residues were performed. The 5E0I crystal structure had missing residues at the spike tops of $\alpha$4 and $\alpha$3 helices for some of the dimers. These residues were modeled based on the other, fully resolved, dimers of these structure. Additionally, because Y132A mutation is located at the inter-dimer interface, the Y132 residue was manually placed in the same position as in the WT 3J2V structure for both 5E0I and 5T2P structures. We used the Molfacture plugin in VMD to construct the different HAP compounds bound to 5E0I structure. The energy of all built compounds was minimized in the binding site, while the protein structure was restrained before addition of water and ions. All systems were solvated and ionized with 0.15 M NaCl, using the solvate and ionize plugins in VMD[1], respectively.

##### MD protocol

All simulations employed the CHARMM36 force field[2] for the capsid protein and the TIP3P model for water[3]. For the drug molecules we use the CGenFF[4] parameters obtained from the PARAMCHEM program[5,6]. Rigid bonds were used for all covalent hydrogen bonds, allowing us to integrate the equations of motion with a 2-fs time step. Van der Waals interactions were cutoff at 12 Å and a smoothing function was applied from 10 to 12 Å to ensure a smooth decay to zero. Long-range electrostatic interactions were calculated using the particle-mesh Ewald method[7]. The temperature and the pressure were kept constant at the biologically relevant values of 310 K and 1 bar, respectively. The Langevin thermostat was used to control the temperature in all simulations. For pressure control we used Langevin barostat[8] in NAMD and Berendsen barostat[9] in AMBER. For the Langevin piston we used a period of 200 fs and a decay of 100 fs[8], while for the Berendsen thermostat we used $\tau$=1.0 ps[9].

##### Definition of base and spike angles

All geometric centers were determined based on the positions of backbone atoms. Base angles were calculated from the centers of $\alpha$5 helices in the tetramer (see Figure S2A). These centers were calculated for each of the two $\alpha$5 helices in a protein dimer, after which the vector going through both centers and pointing away from the tetramer interface was determined. Next, the procedure was repeated for the other dimer in the tetramer, and the base angle was defined as the angle between the two determined vectors. Because the upper parts of the protein spikes (helices $\alpha$4 and $\alpha$3) were highly flexible, we decided to define spike angle based on the lower parts of these helices displayed in Figure S2B. Specifically, the lower spike part was divided into a top and bottom segments; residues 49 to 65 and 103 to 110 defined the top segment, while residues 56 to 65 and 96 to 103 defined the bottom segment. For each dimer we calculated the vector going through the geometric centers of the

the bottom and top segments and pointing towards the spike top. The spike angle was defined as the angle between these two vectors. We investigated if the value of spike angle was sensitive to the exact residue selection by also calculating the same angle using residues 51 to 58 and 101 to 108 for the top segment and residues 58 to 65 and 94 to 101 for the bottom segment. Figure S3 shows that the two different spike angle definitions resulted in very minor differences in the value of this angle.

#### FEP calculations

At each stage of the stratified reaction path, data collection was prefaced by thermalization in the amount of one fourth to one third of the sampling. To augment the accuracy of the free-energy calculation, the alchemical transformations have been performed bidirectionally, and the relative binding free energy was determined using the Bennett acceptance ratio method[10]. Estimation of errors in free-energy calculations is notoriously difficult owing to sources of very different nature, from the finite length of the simulations to force-field inaccuracies[11], presupposing stringent underlying approximations. We have chosen to provide error bars based on the hysteresis between the forward and backward transformations of the bidirectional free-energy calculations, which has proven more realistic than a mere estimate of the statistical error (Table S5). The relative binding free energy between substrates A and B, $\Delta\Delta G(A \rightarrow B)$, is defined as $\Delta G_{bind}(B) - \Delta G_{bind}(A) = \Delta G_{bound} - \Delta G_{unbound}$, where $\Delta G_{bound}$ and $\Delta G_{unbound}$ are the FEP energies for bound and unbound state, respectively. Table S5 shows all calculated FEP energies, while Figure S12 displays the transformations that were used in FEP calculations and the corresponding free energies.

#### Theoretical p$K_a$ calculations

The p$K_a$s of HAP7 and HAP12 were calculated relative to triethylamine (TEA) employing Eq. 1 derived in reference 14[12].

$$pK_{\mathrm{a,HAP}} = pK_{\mathrm{a,TEA}} + 1/(2.303\mathrm{RT})(\Delta\Delta G_{\mathrm{solv,Neu}} - \Delta\Delta G_{\mathrm{solv,H^+}} + \Delta\Delta G_{\mathrm{gas}}) \quad (1)$$

$R$ and $T$ are the ideal gas constant and temperature, respectively. $\Delta\Delta G_{\mathrm{solv,Neu}}$ and $\Delta\Delta G_{\mathrm{solv,H^+}}$ are the differences in solvation free energies between HAP and TEA for the neutral and protonated forms, respectively. Finally, $\Delta\Delta G_{\mathrm{gas}}$ is the difference in gas-phase deprotonation energy between HAP and TEA. All $\Delta\Delta G$ are defined relative to HAP ($\Delta\Delta G = \Delta G_{\mathrm{HAP}} - \Delta G_{\mathrm{TEA}}$). Guassian16 program package[13] was used for calculations of all properties. All geometries were optimized at B3LYP/6-31+G** level of theory. Single point calculations on these geometries were performed with a larger basis set, B3LYP/6-311+G(2d,2p), in both gas phase and water. We used solvation model based on density (SMD) to calculate the solutes solvation free energies in water[14]. Deprotonation energy ($\Delta G_{\mathrm{gas}}$) was calculated as difference in total energies and thermal corrections in gas phase between the deprotonated and protonated state. The total energy was taken from the singe point calculation at B3LYP/6-311+G(2d,2p) level of theory in gas phase. Unfortunately, the HAP compounds were too large for a frequency calculation with this basis set. Therefore, frequencies were calculated at the same level of theory as the geometry optimization and the computed thermal energies were scaled by a factor of 0.977, as suggested for B3LYP/6-31G(d) level of theory[15]. In a previous study a similar approach was shown to have a mean unsigned error of 0.33 p$K_a$ units[12].

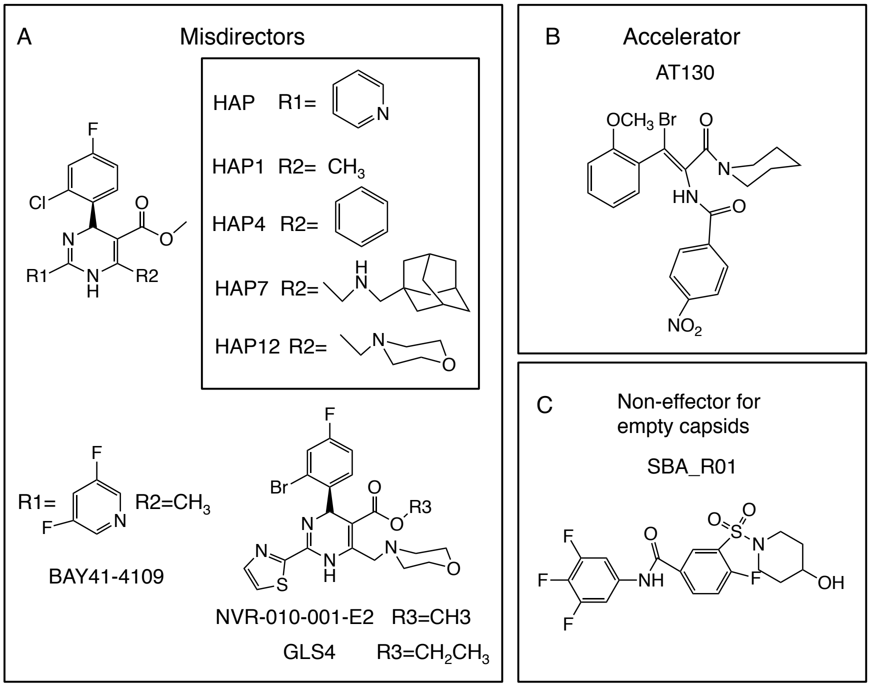


**Figure S1.** Structures of the three known classes of CAMs. A) Heteroaryldihydropyrimidines (HAPs) that misdirect the assembly into non-capsid structures. B) Phenylpropenamides (ATs) that accelerate capsid assembly. C) Sulfamoyl benzamides (SBAs) that do not alter the assembly of empty capsids, yet prevent capsid incorporation of viral DNA.

#### Quasi-Equivalent Structures in HBV capsid

The HBV virus capsid has an icosahedral geometry with triangulation number T = 4[16]. Therefore, it is assembled from 240 copies of the same core protein, HBc, that can assume four different conformations in the shell (A, B, C, or D), depending on its position[17,18]. More specifically, the HBc dimers adopt either AB or CD conformations in the assembled capsid[19,20]. While the two dimers are structurally very similar, larger differences are observed in the quaternary structures for the four quasi-equivalent dimer-dimer contacts found in the capsid. In the assembled capsid there are four quasi-equivalent tetramers (ABCD, DCBA, BAAB, CDCD) and two quasi-equivalent hexamers (CDCDCD and ABCDBA). Note that the dimer interface is not symmetric with respect to the two dimers: the $\alpha 5$ helix of one dimer is wedged between helices $\alpha 5$ and $\alpha 2$ of the other dimer, therefore, ABCD and DCBA are structurally different tetramers. For our tetramer simulations, we chose the ABCD structures, which contain all monomer structures found in the capsid. The structural differences between the four quasi-equivalent tetramers in the crystal structures are relatively small in comparison to the structural fluctuations observed in MD simulations (see Table S1). Additionally, the range of spike and base angles observed in capsid structures for quasi-equivalent tetramers is significantly narrower in comparison to MD simulations (Figure S4). Therefore, we argue that simulating one quasi-equivalent tetramer is sufficient.

For the hexamer, both CDCDCD and ABCDBA hexamers, referred to as symmetric hexamer (SH) and asymmetric hexamer (AH), respectively, were simulated. Three different overlapping tetrameric units are present in the hexamer, resulting in three base and spike angles for each analyzed structure. In AH the different tetramers often showed widely different base and spike angle distributions relative to each other in the same simulation and relative to different simulations (see Figure S5). In contrast, the SH simulations display only slight deviation from the symmetry for the same distributions. Our results suggest that SH is more structurally stable than AH and, therefore, it is a more realistic model for the assembly nucleus. Hence, we have used the results from the simulations of SH for comparison to other structures.

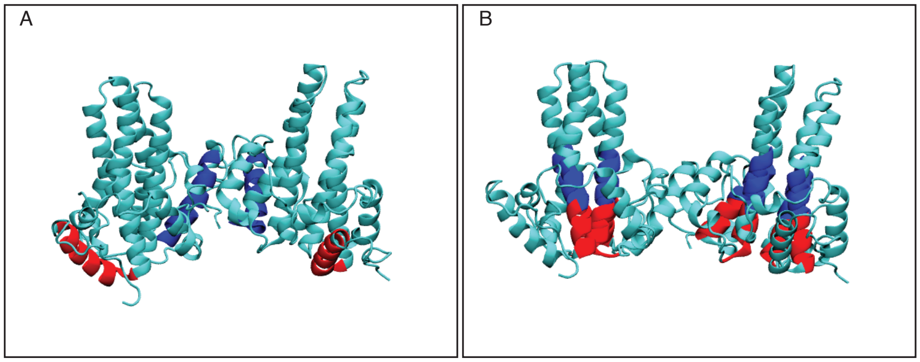


**Figure S2.** Definition of base and spike angles used in our structure analysis. A) $\alpha 5$ helices used for base angle calculations are shown (residues 111 to 127). The outer and inner $\alpha 5$ helices are shown in red and blue, respectively. B) The top and bottom segments of $\alpha 3$ and $\alpha 4$ helices used for spike angle calculation. The top segment is colored blue and the bottom segment is colored red.

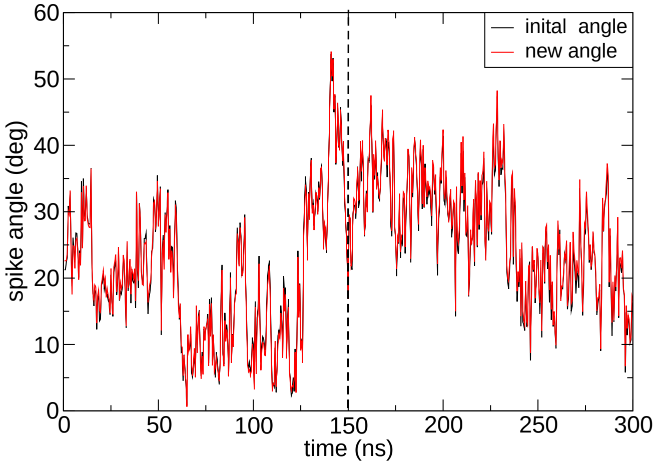


**Figure S3.** Comparison of spike angle values between the initial definition (black) and a slightly altered definition (red) for our apo-tetramer simulation. Dotted line separates the two 150-ns-long simulations.

**Table S1.** Root mean square deviation (RMSD) in Å between the four quasi-equivalent tetramer structures found in assembled capsids. The MD column displays averaged RMSD from the two 150-ns-long MD simulations of the WT ABCD tetramer starting from structure 3J2V relative to each capsid structure.

| Structure | ABCD | DCBA | BAAB | CDCD | MD (300 ns) |
|---|---|---|---|---|---|
| ABCD | 0 | 2.8 | 2.9 | 2.0 | 4.3 |
| CDBA | | 0 | 2.9 | 1.9 | 4.9 |
| BAAB | | | 0 | 1.8 | 5.0 |
| CDCD | | | | 0 | 4.7 |

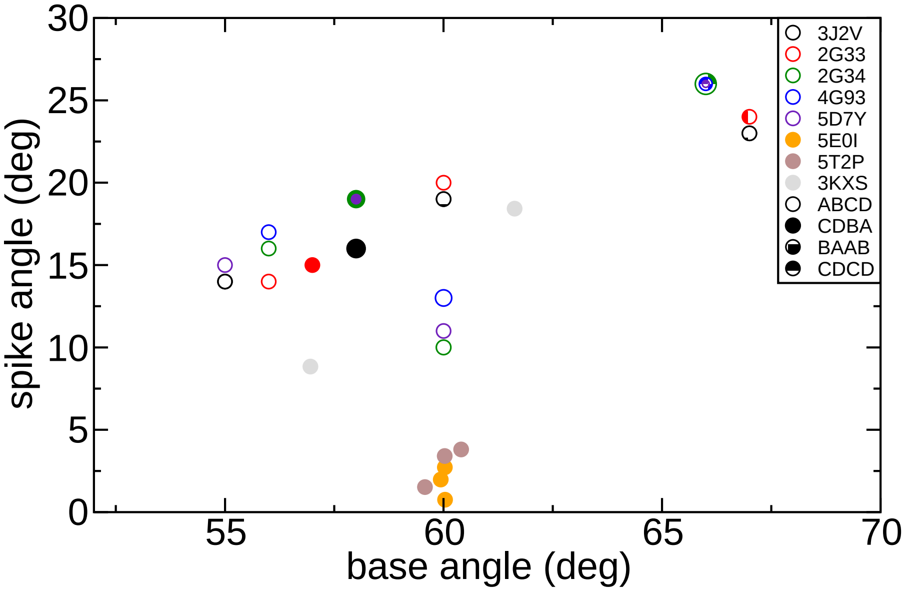


**Figure S4.** Calculated values for spike and base angles for the different quasi-equivalent tetramers in known HBV capsid and hexamer structures. Circle color indicates PDB code, while circle pattern indicates order of the monomers in the tetramer, as shown in the legends. Four quasi-equivalent tetramer structures are found in T4 capsids: ABCD, CDBA, BAAB and CDCD. For hexamer pdb structures (pdb codes 5E0I, 5T2P and 3KXS) the values were calculated for all possible tetramers and plotted in the same color.

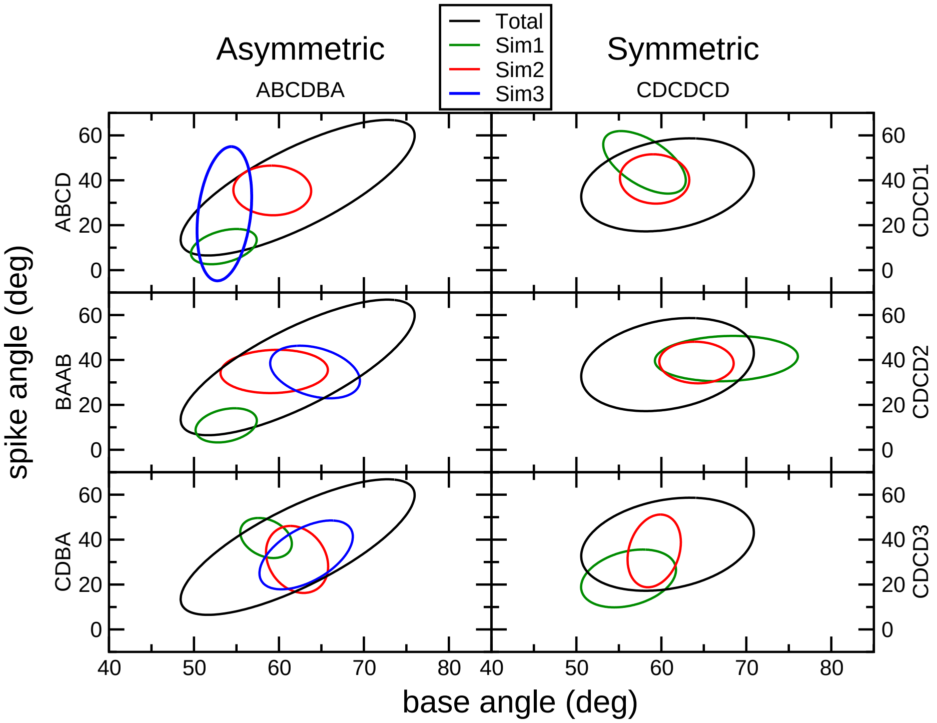


**Figure S5.** Comparison of standard deviation ellipses for different tetramers in the asymmetric (left graphs) and symmetric (right graphs) hexamer. Each graph shows the ellipses for a specific tetramer during different simulations and the ellipse resulting from averages of all tetramers and all simulations (black).

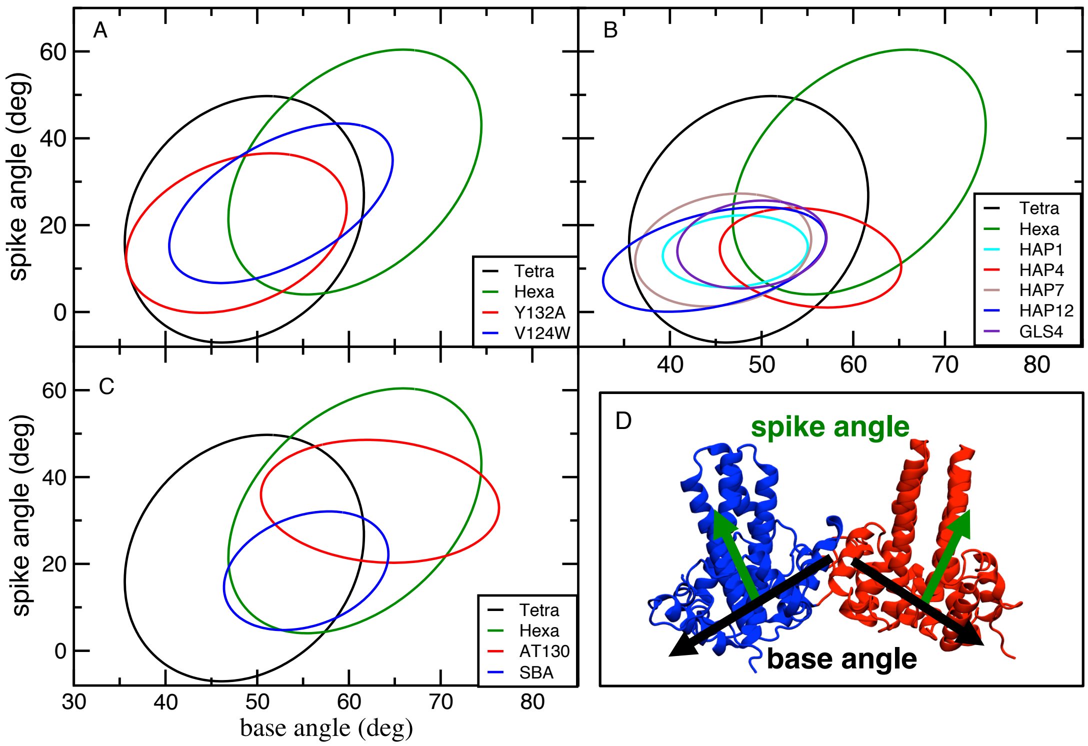


**Figure S6.** Comparison of standard deviation ellipses for apo tetramer and hexamer simulations starting from the hexamer structure (pdb code 5E0I) to the results of mutant and compound-bound tetramers.

#### Repeated simulations

To ensure that our results were not artifacts of the starting structures, we also performed WT tetramer and hexamer simulations using the structure of Y132A mutant hexamer with bound NVR-010-001-E2 (pdb code 5E0I). The drug molecule was removed and the mutation was reversed prior to the simulations. The resulting tetramer simulations showed a similar range of base angles and lower spike angles (see Table S2), in comparison to the simulations of ACBD tetramer from the capsid. There is a close resemblance to the results of Y132A mutant starting from the same structure, suggesting this mutation does not significantly alter tetramer conformations (Figure S6A). One of the two hexamer simulations approximated the results for the SH from the capsid, while the other simulation resembled the results from the AH in terms of base and spike angle sampling (Figure S7). Figure S6 shows that the same conclusions about tetramers, hexamers, mutants and CAMs could be drawn from simulations starting from a different crystal structure.

To further validate our results we also performed simulations of Y132A mutant starting from the capsid ABCD tetramer, and repeated the simulations of the WT tetramer starting from the capsid structure. Figure S8 shows that for both systems one of the simulations agreed with the previous results, while the second simulation sampled base and spike angles found in the hexamer simulations and not in previous apo tetramer simulations. Our simulations suggest that apo-tetramers have a greater flexibility than apo-hexamers or drug-bound tetramers.

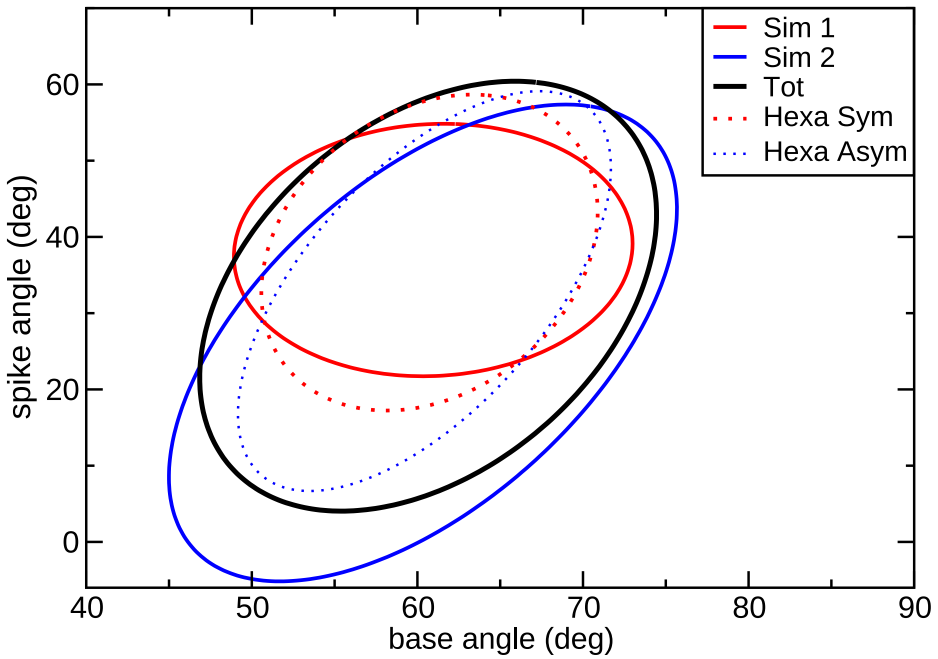


**Figure S7.** Comparison of standard deviation ellipses for the two simulations starting from the hexamer structure 5E0I to the corresponding structures of the symmetric and asymmetric hexamers from 3J2V.

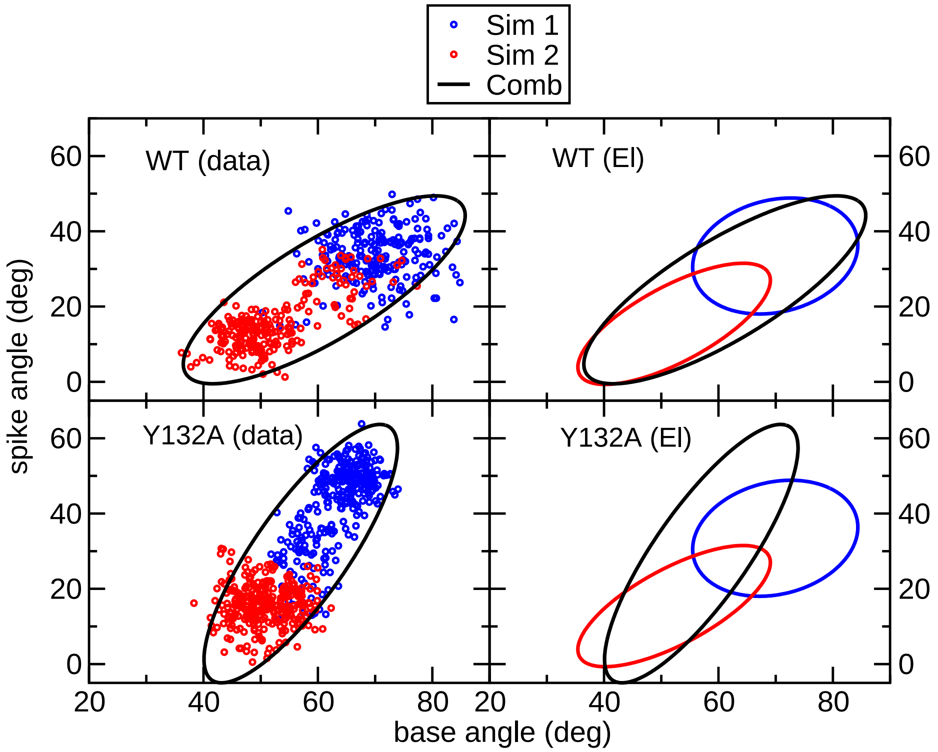


**Figure S8.** Top left: raw data and the resulting standard deviation ellipse for rerun of WT simulation. Top right: comparison of ellipses from the two simulations to the combined ellipse for the rerun of WT simulations. Bottom left: raw data and the resulting standard deviation ellipse for rerun of tetramer with Y132A mutation starting from the capsid state. Bottom right: comparison of ellipses from the two simulations to the combined ellipse for rerun of tetramer with Y132A mutation starting from the capsid state

**Table S2.** The range of sampled base and spike angles in deg for all simulated systems. The values are based on the calculated SDEs.

| System | Base | Spike | System | Base | Spike |
|---|---|---|---|---|---|
| WT Tetra (3J2V) | 31-63 | 1-45 | WT Hexa Asym (3J2V) | 49-72 | 6-59 |
| WT Hexa Sym (3J2V) | 51-71 | 17-59 | Y132A Tetra (3J2V) | 40-74 | -4-65 |
| V124 Tetra (3J2V) | 40-65 | 6-44 | WT Tetra (5E0I) | 36-62 | -7-50 |
| WT Hexa (5E0I) | 47-74 | 4-60 | Y132A Tetra (5E0I) | 35-59 | 1-36 |
| Tetra HAP1 (5E0I) | 39-55 | 6-22 | Tetra HAP4 (5E0I) | 45-65 | 1-24 |
| Tetra HAP7 (5E0I) | 36-55 | 1-27 | Tetra HAP12 (5E0I) | 33-57 | 0-24 |
| Tetra BAY41-4109 (5E0I) | 38-62 | 4-28 | Tetra GLS4 (5E0I) | 41-57 | 5-26 |
| Hexa 1 GLS4 (5E0I) | 55-66 | -2 -23 | Hexa 3 GLS4 (5E0I) | 57-63 | -1-17 |
| Tetra AT130 (4G93) | 50-76 | 20-49 | Hexa 1 AT130 (4G93) | 56-66 | 26-49 |
| Hexa 3 AT130 (4G93) | 52-69 | 22-47 | Tetra SBA_R01 (5T2P) | 46-64 | 5-32 |
| Tetra GT-5 (Docking) | 45-66 | 17-50 | Tetra GT-39 (Docking) | 37-61 | -1-27 |
| Tetra GT-40 (Docking) | 43-60 | 3-30 | Tetra GT-45 (Docking) | 45-63 | 3-30 |
| Tetra GT-46 (Docking) | 45-65 | 6-32 | Tetra GT-47 (Docking) | 44-66 | 13-40 |

#### Additional simulation data

**Table S3.** Average values (Av) and standard errors (SE) in deg for the studied systems. SEs were calculated from averages of the two independent simulations.

| System | Base Angle | | Spike Angle | |
|---|---|---|---|---|
| | Av | SE | Av | SE |
| GT39 | 49.4 | 2.0 | 12.9 | 1.4 |
| GT40 | 51.5 | 0.8 | 16.9 | 1.9 |
| GT45 | 53.9 | 2.5 | 15.9 | 2.5 |
| GT46 | 55.0 | 0.2 | 19.3 | 3.2 |
| GT47 | 54.9 | 0.6 | 26.5 | 2.4 |
| GT5 | 55.5 | 2.7 | 33.2 | 0.3 |
| 1 AT130 Hexa | 60.7 | $<0.1$ | 37.6 | 0.2 |
| 1 GLS4 Hexa | 60.1 | $<0.1$ | 10.4 | 1.8 |
| 3 AT130 Hexa | 60.7 | 0.1 | 34.4 | 0.3 |
| 3 GLS4 Hexa | 60.1 | $<0.1$ | 8.1 | 0.4 |
| Y132A (3J2V) | 57.1 | 6.6 | 30.1 | 14.3 |
| AT130 | 63.4 | 3.0 | 34.6 | 2.1 |
| BAY | 50.0 | 2.2 | 15.9 | 0.2 |
| GSL4 | 49.0 | 0.4 | 15.5 | 0.6 |
| HAP1 | 47.1 | 1.0 | 14.0 | 0.5 |
| HAP12 | 44.9 | 1.9 | 12.1 | 2.9 |
| HAP4 | 55.3 | 1.2 | 12.5 | 2.1 |
| HAP7 | 45.8 | 2.6 | 14.3 | 2.3 |
| Hexa Asym | 62.2 | 1.6 | 36.7 | 2.6 |
| Hexa Sym | 60.7 | 0.1 | 37.9 | 0.6 |
| SBA | 55.3 | 0.4 | 18.4 | 0.7 |
| V124W | 52.4 | 3.9 | 25.2 | 5.1 |
| Y132A (5E0I) | 47.4 | 0.8 | 18.9 | 2.7 |
| WT Tetra (3J2V) | 47.0 | 4.4 | 23.2 | 3.5 |
| WT Tetra (5E0I) | 48.6 | 1.2 | 21.3 | 7.3 |

| | AT130 Hexa | GLS4 Hexa | AT130 Tetra | GSL4 Tetra | GT | | | | | HAP | | | | Hexa | SBA | V124W | Y132A | Tetra |
|---|---|---|---|---|---|---|---|---|---|---|---|---|---|---|---|---|---|---|
| | | | | | 39 | 45 | 46 | 47 | 5 | 1 | 12 | 4 | 7 | | | | | |
| AT130 (Hexa) | **0.9** | 0.0 | 0.97 | 0.0 | 0.0 | 0.01 | 0.12 | 0.41 | 0.95 | 0.0 | 0.0 | 0.0 | 0.0 | 1.0 | 0.12 | 0.7 | 0.03 | 0.0 |
| GLS4 (Hexa) | 0.0 | **0.83** | 0.01 | 0.06 | 0.47 | 0.49 | 0.51 | 0.27 | 0.13 | 0.0 | 0.06 | 0.8 | 0.0 | 0.13 | 0.5 | 0.13 | 0.08 | 0.65 |
| AT130 Tetra | 0.31 | 0.0 | **0.75** | 0.0 | 0.0 | 0.08 | 0.16 | 0.35 | 0.56 | 0.0 | 0.0 | 0.0 | 0.0 | 0.76 | 0.15 | 0.42 | 0.13 | 0.11 |
| GSL4 Tetra | 0.0 | 0.05 | 0.0 | **0.91** | 1.0 | 0.74 | 0.69 | 0.4 | 0.23 | 0.72 | 0.94 | 0.63 | 0.91 | 0.08 | 0.62 | 0.91 | 0.98 | 1.0 |
| GT39 | 0.0 | 0.19 | 0.0 | 0.48 | **0.76** | 0.58 | 0.5 | 0.29 | 0.17 | 0.38 | 0.64 | 0.53 | 0.67 | 0.09 | 0.48 | 0.55 | 0.73 | 0.83 |
| GT45 | 0.01 | 0.28 | 0.12 | 0.51 | 0.83 | **0.66** | 0.85 | 0.53 | 0.37 | 0.35 | 0.5 | 0.73 | 0.48 | 0.29 | 0.86 | 0.66 | 0.65 | 0.94 |
| GT46 | 0.05 | 0.27 | 0.22 | 0.44 | 0.66 | 0.77 | **0.75** | 0.68 | 0.5 | 0.28 | 0.4 | 0.6 | 0.4 | 0.39 | 0.87 | 0.72 | 0.64 | 0.84 |
| GT47 | 0.16 | 0.13 | 0.43 | 0.23 | 0.35 | 0.45 | 0.62 | **0.83** | 0.8 | 0.11 | 0.2 | 0.28 | 0.22 | 0.59 | 0.55 | 0.85 | 0.61 | 0.68 |
| GT5 | 0.3 | 0.05 | 0.56 | 0.1 | 0.17 | 0.25 | 0.37 | 0.64 | **0.75** | 0.03 | 0.08 | 0.12 | 0.1 | 0.71 | 0.33 | 0.69 | 0.39 | 0.45 |
| HAP1 | 0.0 | 0.0 | 0.0 | 0.91 | 1.0 | 0.64 | 0.56 | 0.24 | 0.08 | **0.87** | 1.0 | 0.56 | 1.0 | 0.0 | 0.51 | 0.91 | 1.0 | 0.96 |
| HAP12 | 0.0 | 0.03 | 0.0 | 0.57 | 0.82 | 0.45 | 0.39 | 0.21 | 0.11 | 0.49 | **0.76** | 0.38 | 0.81 | 0.04 | 0.36 | 0.56 | 0.86 | 0.67 |
| HAP4 | 0.0 | 0.48 | 0.0 | 0.46 | 0.8 | 0.77 | 0.7 | 0.35 | 0.19 | 0.33 | 0.46 | **0.77** | 0.41 | 0.12 | 0.71 | 0.47 | 0.48 | 0.9 |
| HAP7 | 0.0 | 0.0 | 0.0 | 0.61 | 0.92 | 0.46 | 0.42 | 0.26 | 0.14 | 0.53 | 0.88 | 0.37 | **0.69** | 0.03 | 0.37 | 0.66 | 0.98 | 0.8 |
| Hexa | 0.27 | 0.04 | 0.66 | 0.03 | 0.08 | 0.17 | 0.25 | 0.41 | 0.61 | 0.0 | 0.02 | 0.07 | 0.02 | **0.7** | 0.23 | 0.45 | 0.2 | 0.2 |
| SBA | 0.06 | 0.28 | 0.22 | 0.43 | 0.69 | 0.86 | 0.95 | 0.66 | 0.49 | 0.28 | 0.41 | 0.67 | 0.38 | 0.4 | **0.87** | 0.69 | 0.61 | 0.85 |
| V124W | 0.2 | 0.05 | 0.38 | 0.39 | 0.49 | 0.41 | 0.49 | 0.62 | 0.64 | 0.31 | 0.39 | 0.27 | 0.43 | 0.48 | 0.42 | **0.6** | 0.71 | 0.72 |
| Y132A (5E0I) | 0.01 | 0.03 | 0.12 | 0.42 | 0.65 | 0.4 | 0.44 | 0.45 | 0.36 | 0.34 | 0.61 | 0.28 | 0.63 | 0.21 | 0.38 | 0.72 | **0.65** | 0.82 |
| Tetra | 0.0 | 0.15 | 0.07 | 0.27 | 0.47 | 0.37 | 0.36 | 0.32 | 0.26 | 0.21 | 0.3 | 0.33 | 0.32 | 0.14 | 0.33 | 0.45 | 0.52 | **0.72** |

**Table S4.** Fractional area overlaps for SDEs. Each row shows the fractional overlap area of the total SDE for that system with the systems specified in each column. To estimate the differences between the two independent simulations in each system, we calculated the overlap area between the SDE of each independent simulation and the SDE from the two combined simulations as a fraction of the latter SDE area. Averages of these fractional areas from the two independent simulations are shown as diagonal values in bold. Because there is no mathematical formula to calculate overlap area between two ellipses we used numerical integration to calculate all overlap areas.

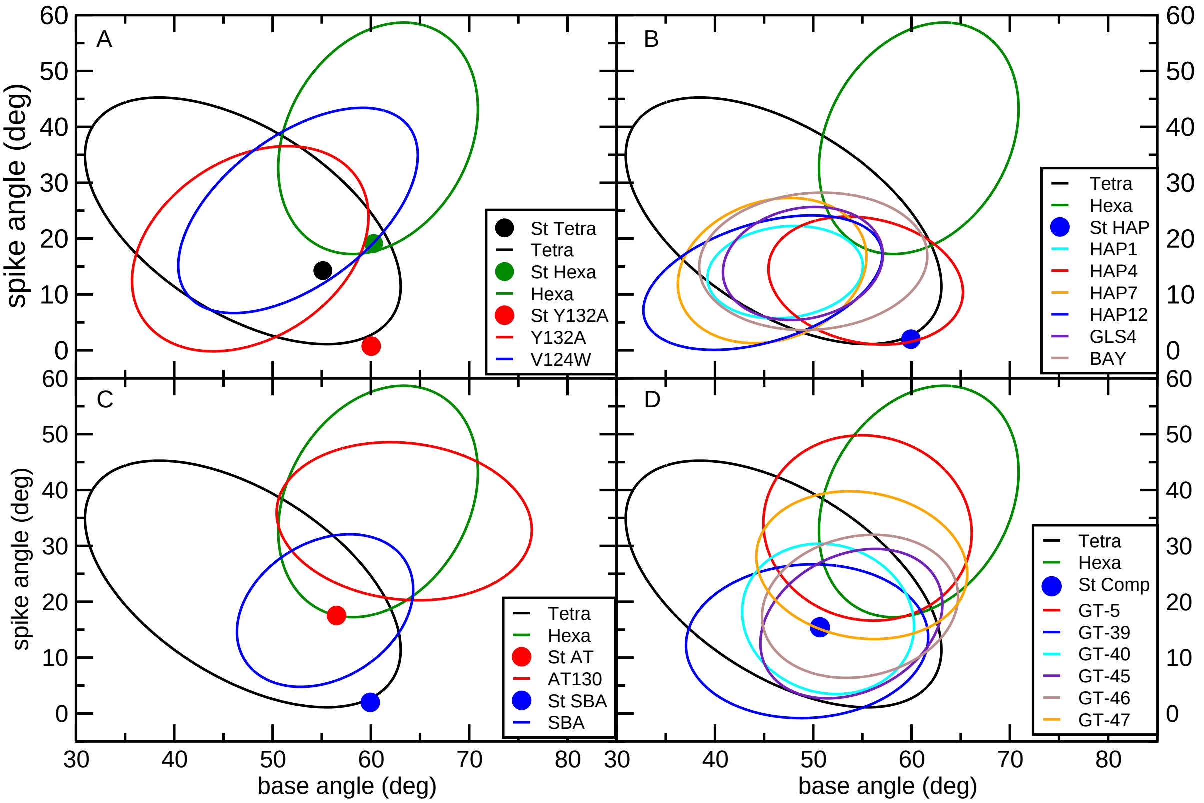


**Figure S9.** Standard deviation ellipses for all simulated tetramers. Symmetric hexamer results are added for comparison. In addition, the spike and base values for starting structures are also shown.

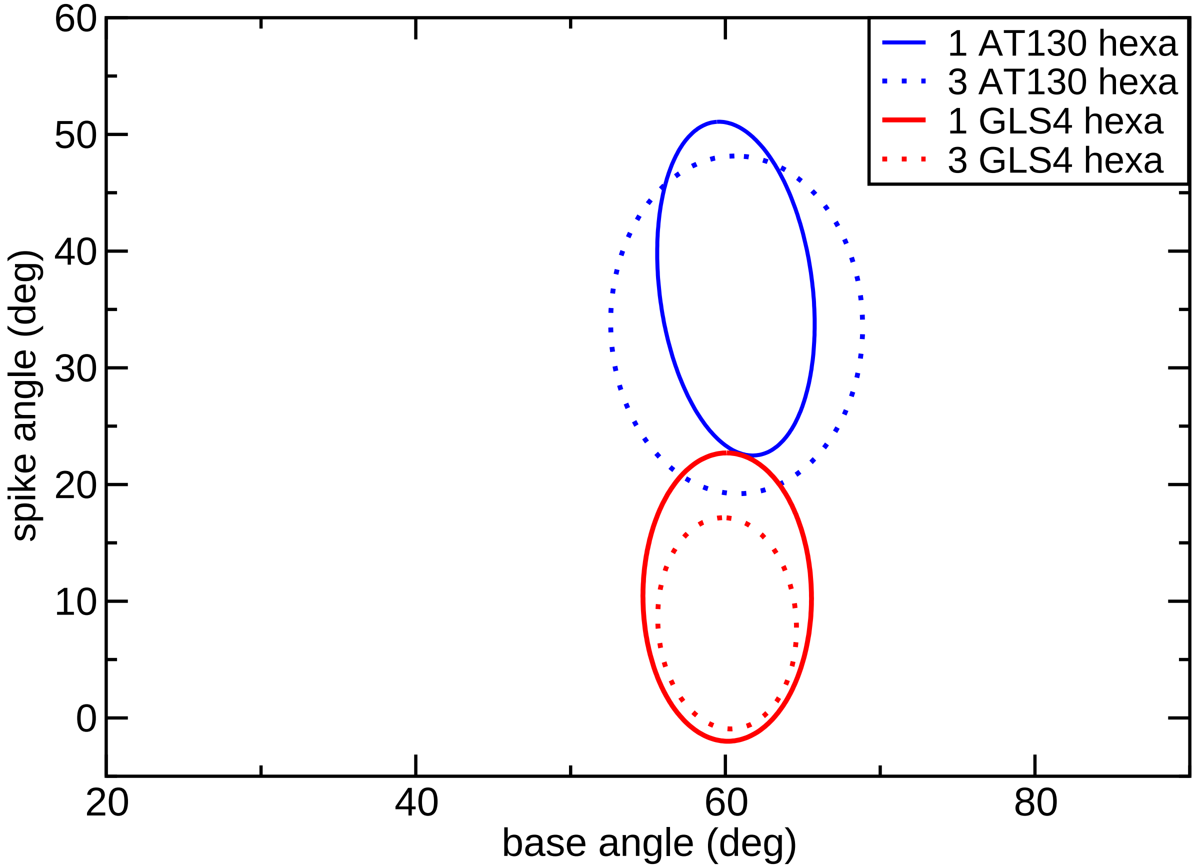


**Figure S10.** Standard deviation ellipses for AT130 and GLS4 with either one of three (one at each tetrameric interface) bound molecules. FOAs between one molecule and three molecule systems were 94% and 60%, for AT130 and GLS4, respectively.

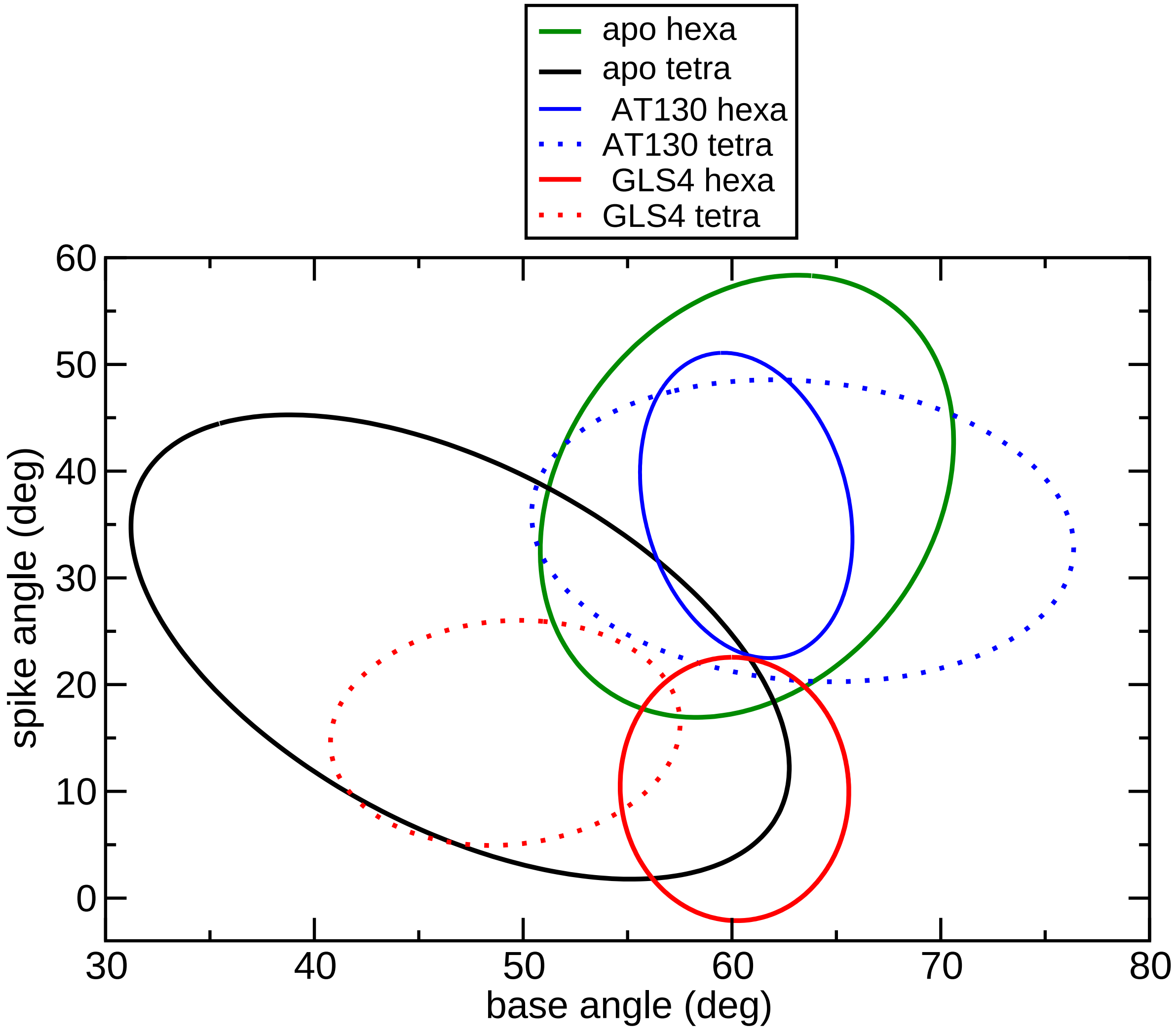


**Figure S11.** Standard deviation ellipses for hexamers with bound one GLS4 or AT130. The results for apo tetramer and hexamer, as well as tetramers with bound AT130 and GLS4, are added for comparison.

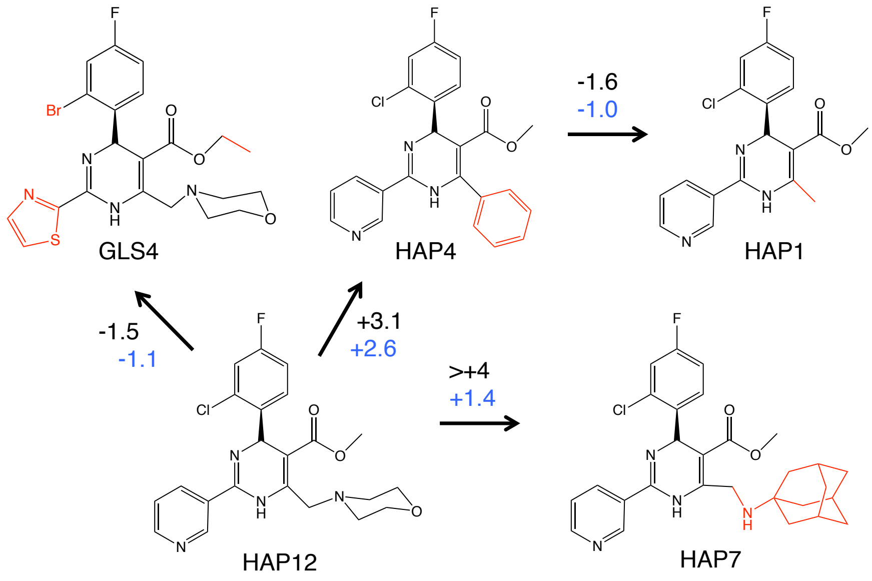


**Figure S12.** Setup and results of our FEP calculations. The structural differences from HAP12 are shown in red. All differences in binding free energies are shown in kcal/mol. Black values were calculated from experimental $EC_{50}$s using Eq. 1 in the main text, while blue values were obtained from FEP calculations.

**Table S5.** Summary of the alchemical free-energy calculations performed to estimate the relative binding free energies of HAP1, HAP4, HAP7 and GLS4 to Cp149 tetramer. $\Delta G_{forward}$ and $\Delta G_{backward}$ represent the free-energy perturbation estimates for the annihilation and creation of the substrate in the bound and in the unbound states. $\Delta G_{BAR}$ is the Bennett acceptance ratio estimate of the binding free energy, $\Delta G_{bind}$, based on the bidirectional transformations[10]. All free energies are given in kcal/mol.

| | time (ns) | $\Delta G_{forward}$ | $\Delta G_{backward}$ | $\Delta G_{BAR}$ | $\Delta\Delta G_{exp}$ |
|---|---|---|---|---|---|
| HAP12 → HAP7 | | | | | |
| bound | 80 | +68.1 | -64.6 | +66.0±1.7 | |
| unbound | 40 | +64.4 | -64.8 | +64.6±0.2 | |
| ΔΔG | | | | +1.4±1.9 | > 4 |
| HAP12 → HAP4 | | | | | |
| bound | 15 | -11.0 | +11.1 | -11.3±0.0 | |
| unbound | 15 | -13.7 | +14.1 | -13.9±0.2 | |
| ΔΔG | | | | +2.6±0.2 | +3 |
| HAP12 → GLS4 | | | | | |
| bound | 15 | -23.2 | +18.4 | -20.8±2.4 | |
| unbound | 15 | -19.4 | +20.0 | -19.7±0.3 | |
| ΔΔG | | | | -1.1±2.4 | -0.5 |
| HAP4 →HAP1 | | | | | |
| bound | 15 | -26.2 | +28.5 | -27.5±1.2 | |
| unbound | 15 | -26.3 | +26.3 | -26.5±0.0 | |
| ΔΔG | | | | -1.0±1.2 | -1.6 |

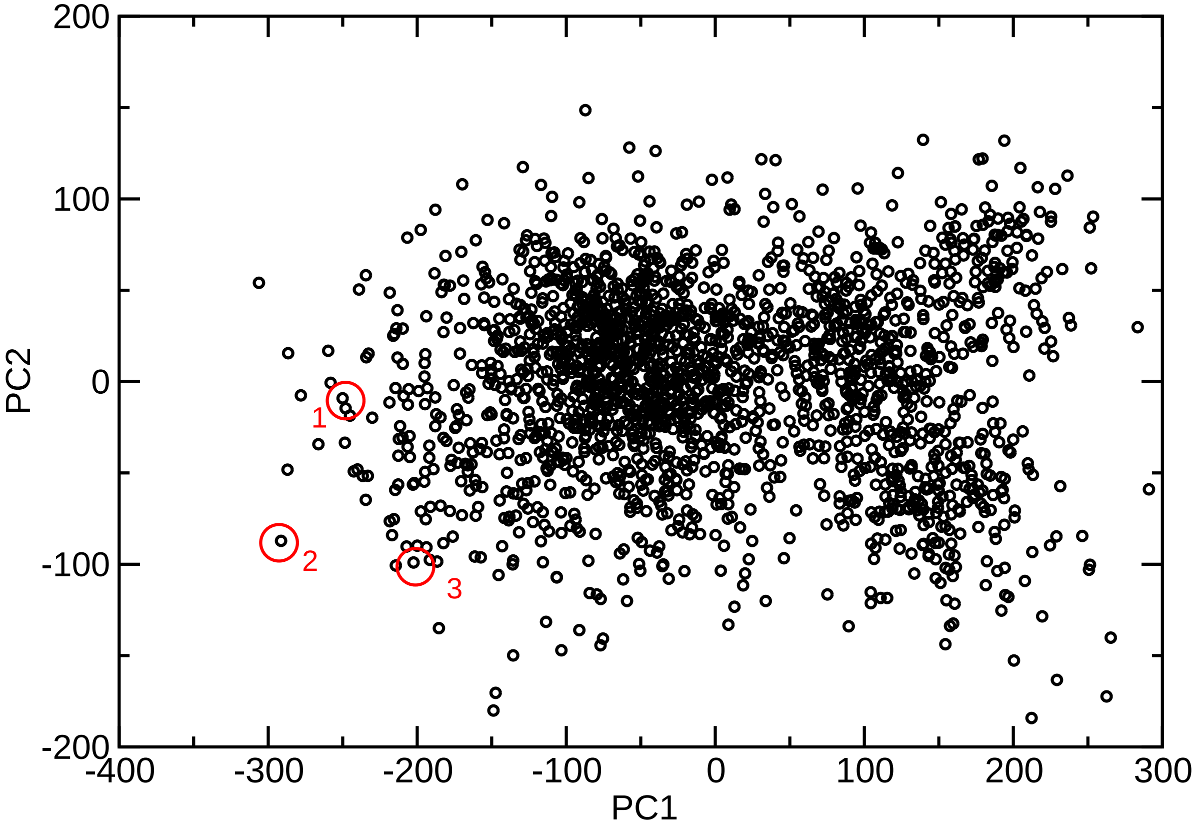


**Figure S13.** 2D plot of the two largest principal components in our PCA analysis of apo tetramer. The structures that were selected for docking are marked in red.

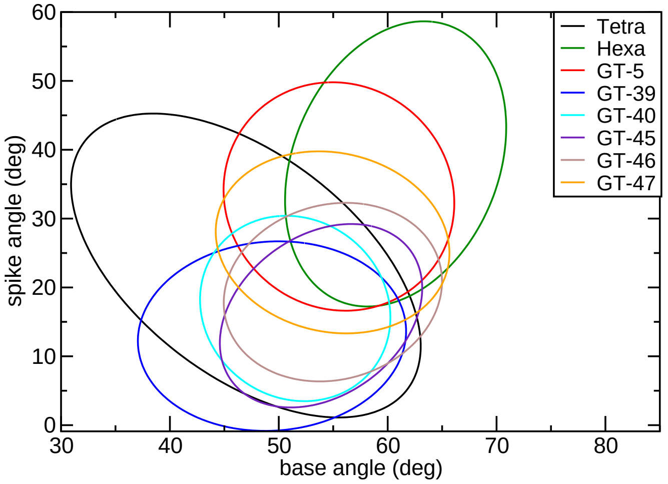


**Figure S14.** SDEs of base and spike angles for all of our novel compounds. Apo tetramer and hexamer SDEs are added for comparison.

**Table S6.** Summarized results of HBV DNA inhibition and toxicity testing for the first round of selected compounds. N/a stands for not applied.

| Comp. | HBV DNA inhibition (%) at 10 $\mu$M | MTT cytotoxicity, $IC_{50}$ ($\mu$M) | | | |
|---|---|---|---|---|---|
| | | PBM | CEM | Vero | HepG2 |
| 1 | <1 | 82.9 | 26.9 | >100 | 12 |
| 2 | <1 | 85.3 | 16.1 | >100 | 26.4 |
| 3 | 40.4 | 34.0 | 15.4 | 27.0 | 51.2 |
| 4 | <1 | >100 | >100 | >100 | >100 |
| 5 | 24.7 | >100 | >100 | >100 | >100 |
| 6 | 27.2 | 47.7 | 18.7 | 42.6 | 14.9 |
| 7 | <1 | >100 | >100 | >100 | >100 |
| 8 | <1 | >100 | >100 | >100 | >100 |
| 9 | 34.6 | >100 | >100 | >100 | 55.3 |
| 10 | <1 | 18.2 | 15.2 | 14.4 | 12.9 |
| 11 | <1 | 63.9 | 13.3 | 18.9 | 31.6 |
| 12 | <1 | >100 | 17.0 | >100 | >100 |
| 13 | <1 | >100 | >100 | >100 | >100 |
| 14 | <1 | >100 | >100 | >100 | >100 |
| 15 | <1 | >100 | >100 | >100 | >100 |
| 16 | 1.7 | >100 | >100 | >100 | >100 |
| 17 | <1 | >100 | >100 | >100 | >100 |
| 18 | <1 | >100 | >100 | >100 | >100 |
| 19 | <1 | >100 | >100 | >100 | >100 |
| 20 | <1 | >100 | >100 | >100 | >100 |
| 21 | <1 | >100 | 18.2 | 49.6 | >100 |
| 22 | <1 | >100 | 36.3 | 88.6 | >100 |
| 23 | <1 | >100 | >100 | >100 | >100 |
| 24 | <1 | >100 | 11.3 | >100 | >100 |
| 25 | <1 | >100 | >100 | >100 | >100 |
| 26 | <1 | >100 | >100 | >100 | >100 |
| 27 | <1 | 14.7 | 4.0 | 27.3 | 41.7 |
| 28 | <1 | 24.0 | 63.8 | >100 | >100 |
| 29 | <1 | 25.0 | 35.7 | 21.0 | 12.3 |
| 3TC | 97 | 42.4 | 22.5 | >100 | >100 |
| Cycloheximide | n/a | 0.9 | 0.2 | 0.2 | 0.3 |

#### Structures and experimental data for all tested compounds

**1**

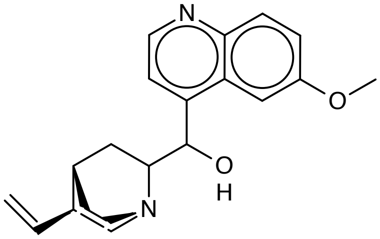


**2**

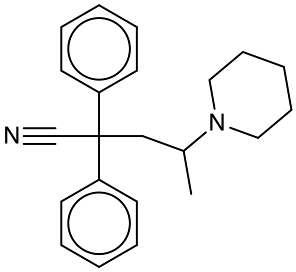


**3**

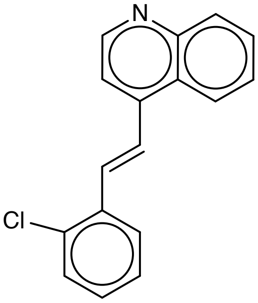


**4**

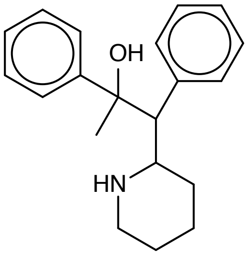


**5**

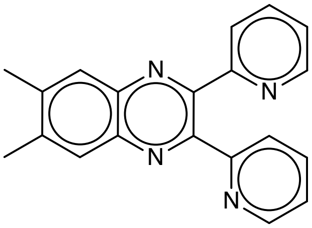


**6**

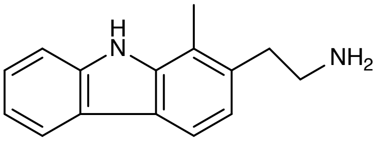


**7**

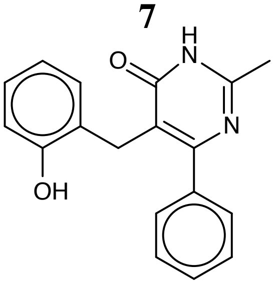


**8**

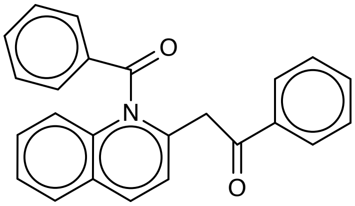


**9**

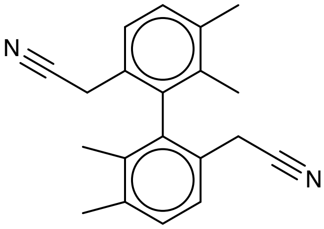


**10**

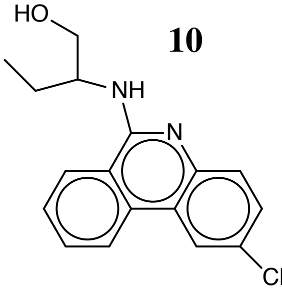


**11**

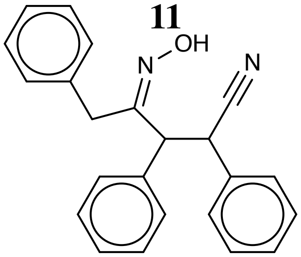


**Figure S15.** Structures of compounds 1-11.

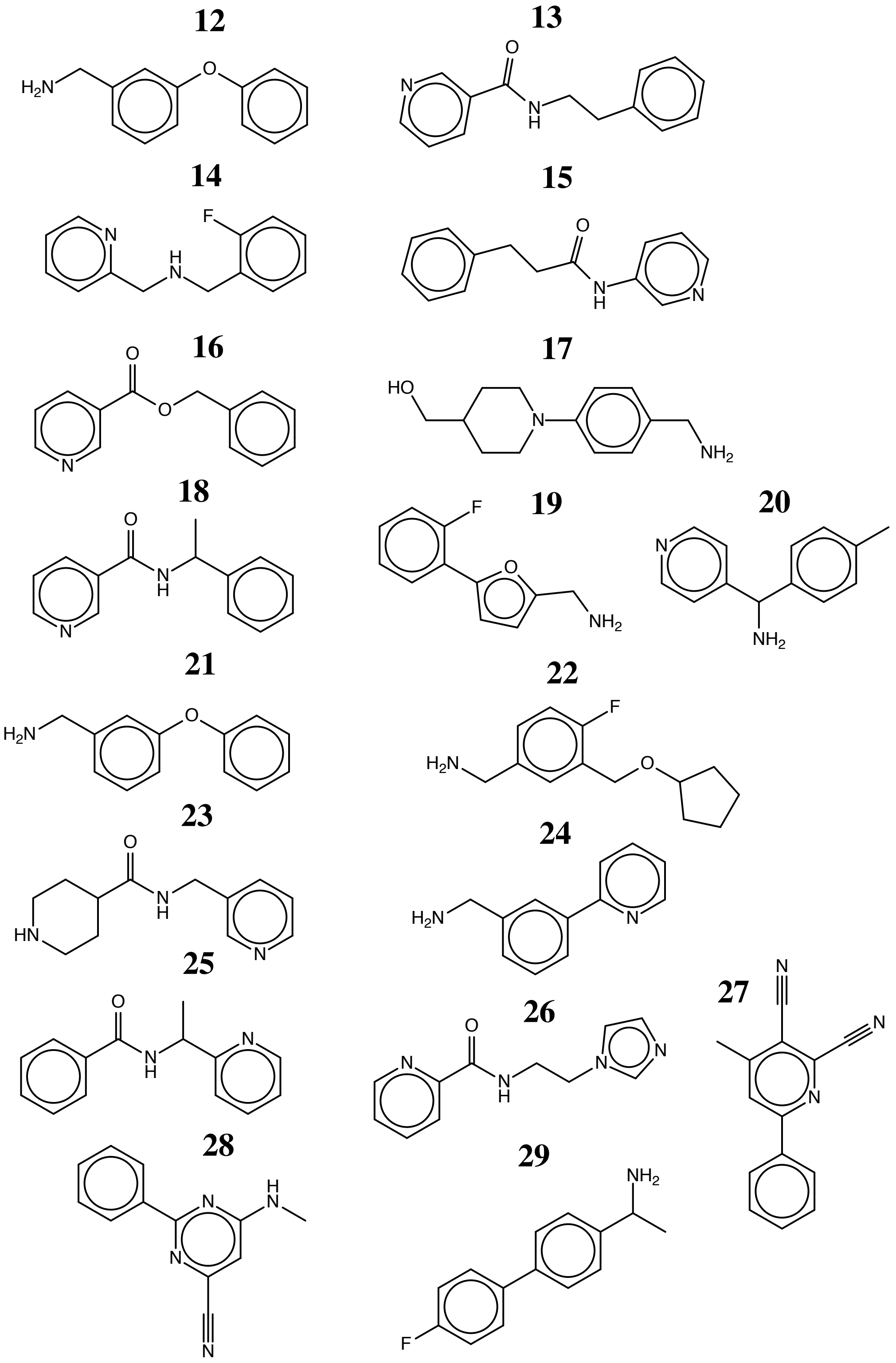


**Figure S16.** Structures of compounds 12-29.

**30**

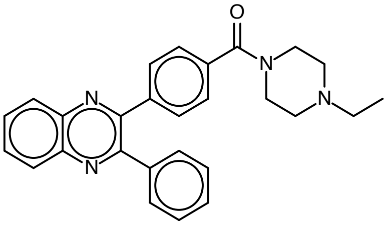

**31**

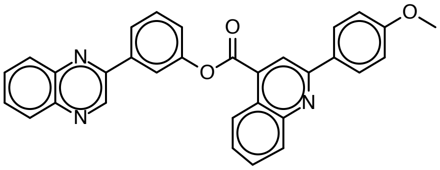

**32**

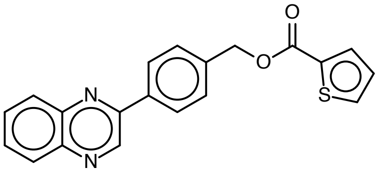

**33**

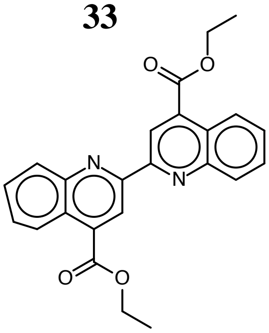

**34**

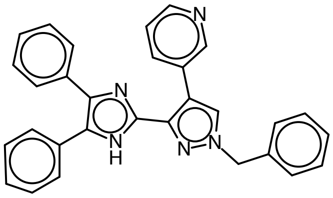

**35**

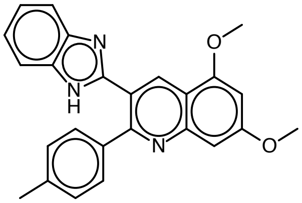

**36**

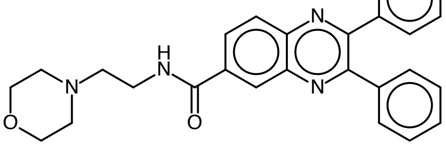

**37**

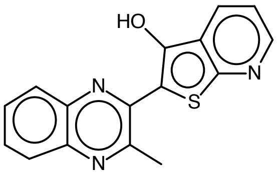

**38**

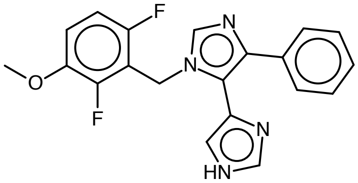

**39**

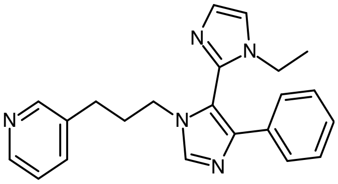

**40**

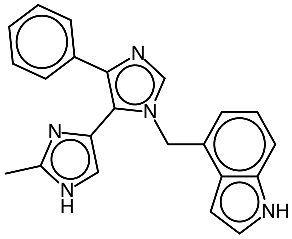

**41**

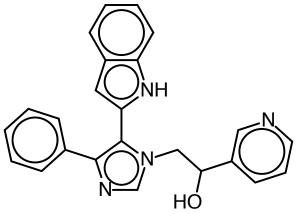

**Figure S17.** Structures of compounds 30-41.

**42**

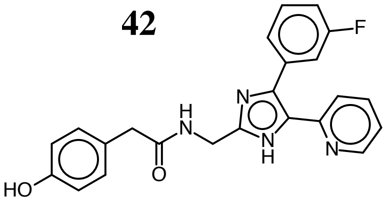


**43**

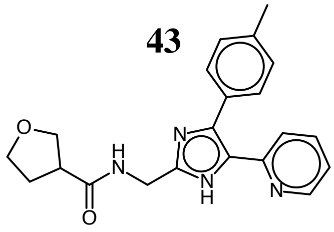


**44**

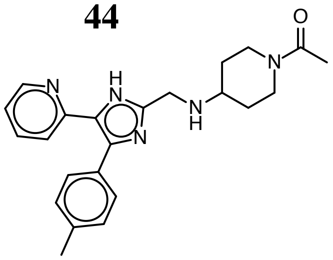


**45**

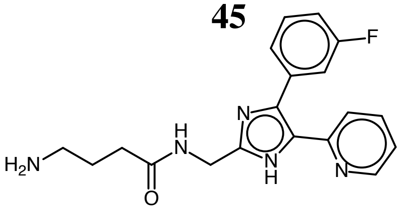


**46**

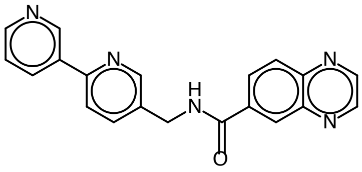


**47**

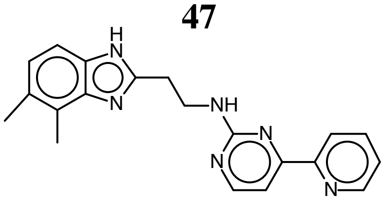


**48**

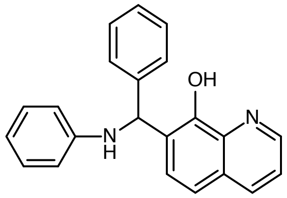


**49**

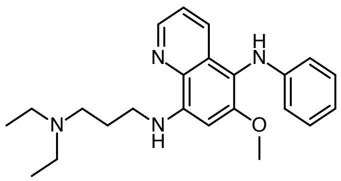


**Figure S18.** Structures of compounds 42-49.

**Table S7.** Summarized results of HBV DNA inhibition and toxicity testing for the second round of selected compounds.

| Comp. | Inhibition | MTT cytotoxicity, $IC_50$ ($\mu$M) (inhibition at 100 $\mu$M) | | | |
|---|---|---|---|---|---|
| | (%) at 10 $\mu$M | PBM | CEM | Vero | HepG2 |
| 30 | <1 | 60 | 32 | 81 | 34 |
| 31 | <1 | 86 | >100 (36%) | 97 | 42 |
| 32 | 48 | 85 | 52 | 44 | 59 |
| 33 | 60 | >100 (5.6%) | >100 (32%) | 76 | 86 |
| 34 | <1 | 59 | 11 | 14 | 16 |
| 35 | 36 | >100 (14%) | >100 (49%) | 42 | 63 |
| 36 | 11 | 10 | 18 | 31 | 29 |
| 37 | 40 | >100 (48%) | >100 (<1%) | 19 | 24 |
| 38 | 45 | >100 (24%) | 62 | 48 | 24 |
| 39 | 57 | >100 (22%) | 16 | >100 (47%) | >100 (32%) |
| 40 | 55 | >100 (19%) | 77 | >100 (39%) | >100 (30%) |
| 41 | <1 | 27 | 12 | 18 | 10 |
| 42 | <1 | 3.5 | 1.4 | 13 | 5.4 |
| 43 | 4.7 | 42 | 30 | 19 | 29 |
| 44 | 37 | 64 | 63 | >100 (54%) | 70 |
| 45 | 50 | >100 (41%) | >100 (36%) | 66 | >100 (41%) |
| 46 | 49 | >100 (20%) | 38 | 53 | 82 |
| 47 | 49 | >100 (16%) | >100 (48%) | >100 (31%) | 18 |
| 48 | <1 | 20 | 3.4 | 4.5 | 34 |
| 49 | 45 | 27 | 9.4 | 16 | 17 |
| 3TC | 82 | 42 | 22 | >100 | >100 |

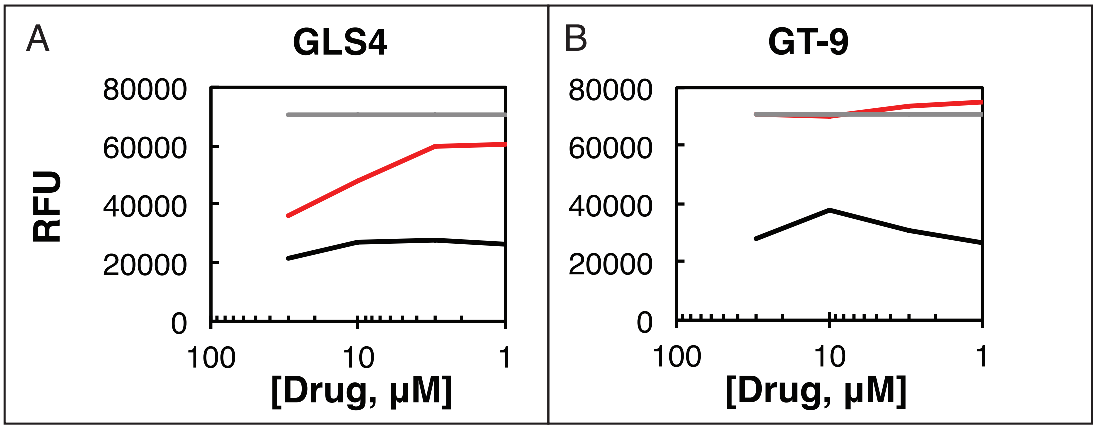


**Figure S19.** Measurement of HBV Cp149 tryptophan fluorescence in relative fluorescence units (RFU) in the presence of GLS4 and GT-9 (A and B, respectively). The grey line shows data for HBV Cp149 dimer only in DMSO, the black line shows compound only in buffer, and the red line shows CAM in presence on Cp149 dimer.

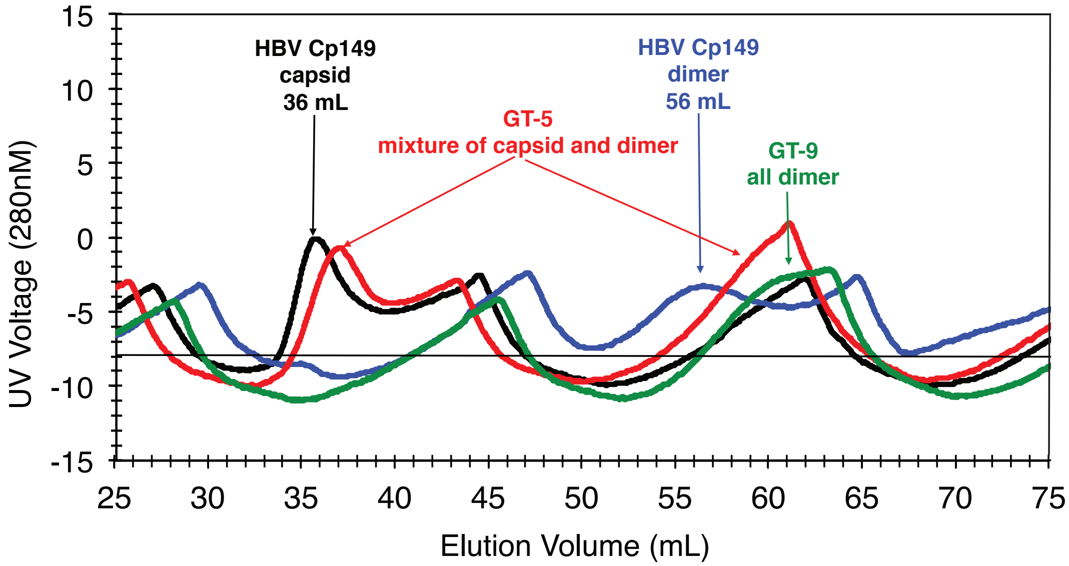


**Figure S20.** The results of analytical size-exclusion chromatography of Cp149 with and without our compounds. The black line shows assembled capsid, the blue line shows Cp149 dimer, and finally the red and green lines show Cp149 dimer in presence of GT-5 and GT-9, respectively.

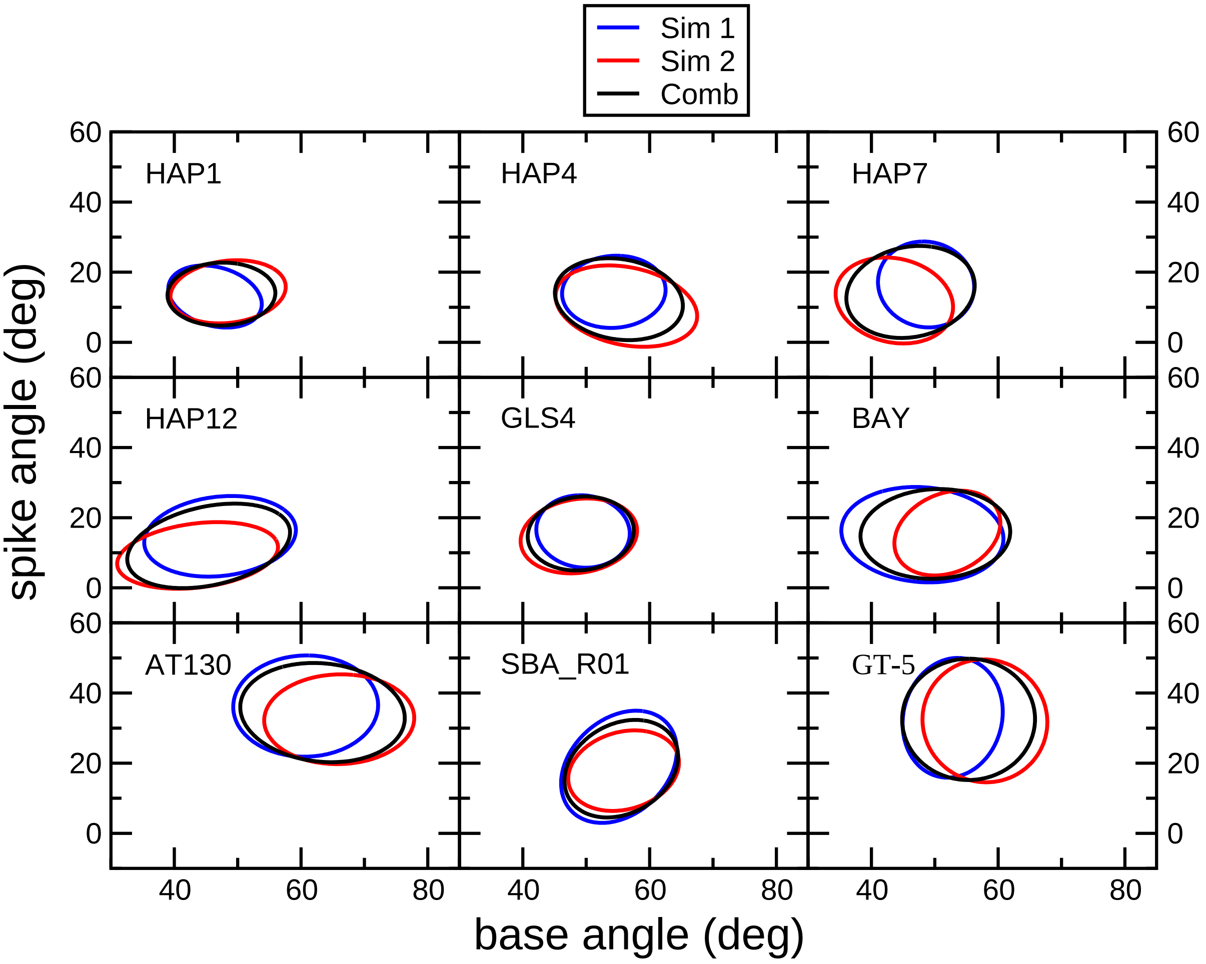


**Figure S21.** Comparison of standard deviation ellipses resulting from each of the two simulations to the one obtained from combining both simulations for the studied tetramers with bound drug compounds.

#### Convergence of simulations

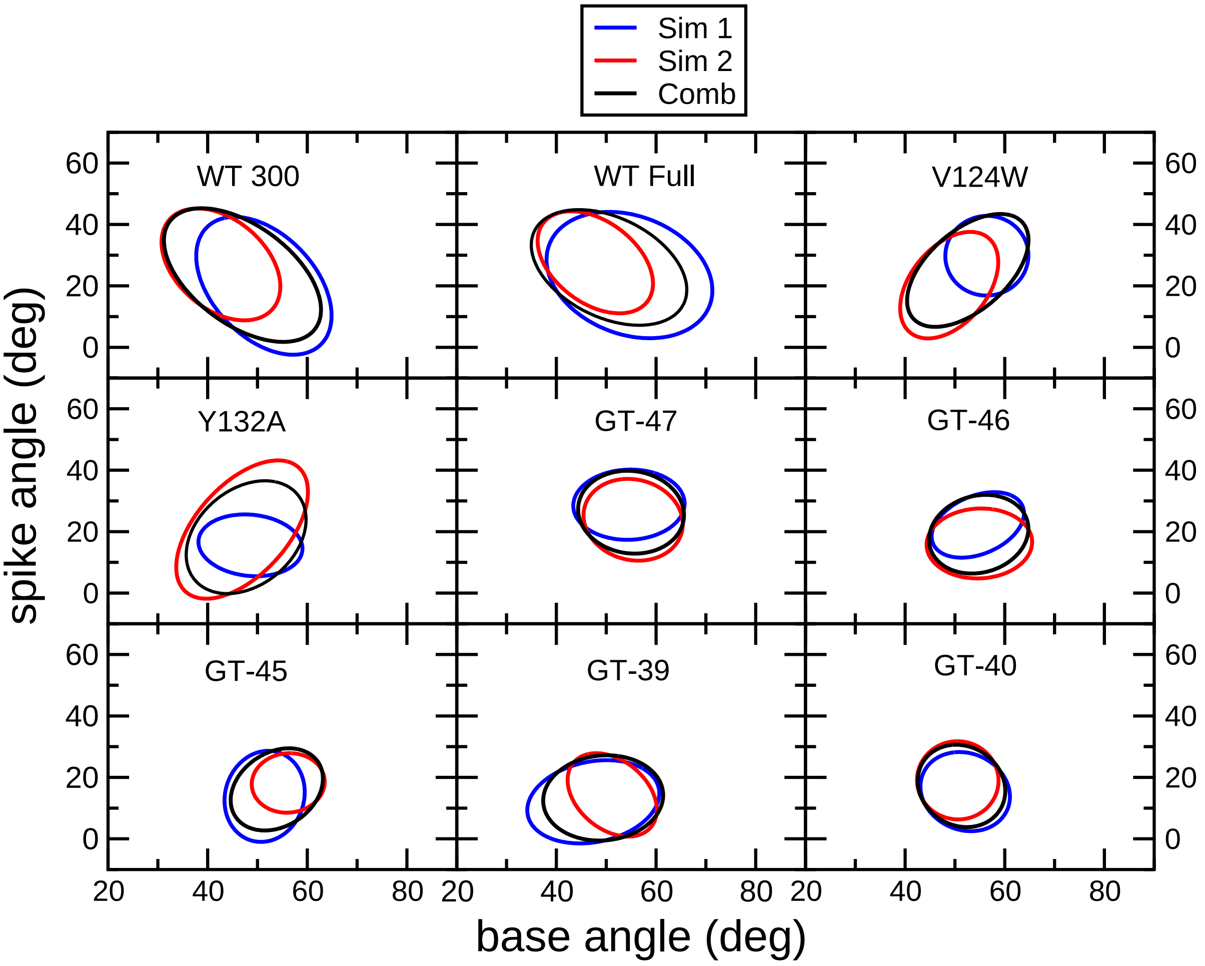


**Figure S22.** Comparison of standard deviation ellipses resulting from each of the two simulations to the one obtained from combining both simulations for the studied apo-tetramers and tetramers with bound novel compounds.

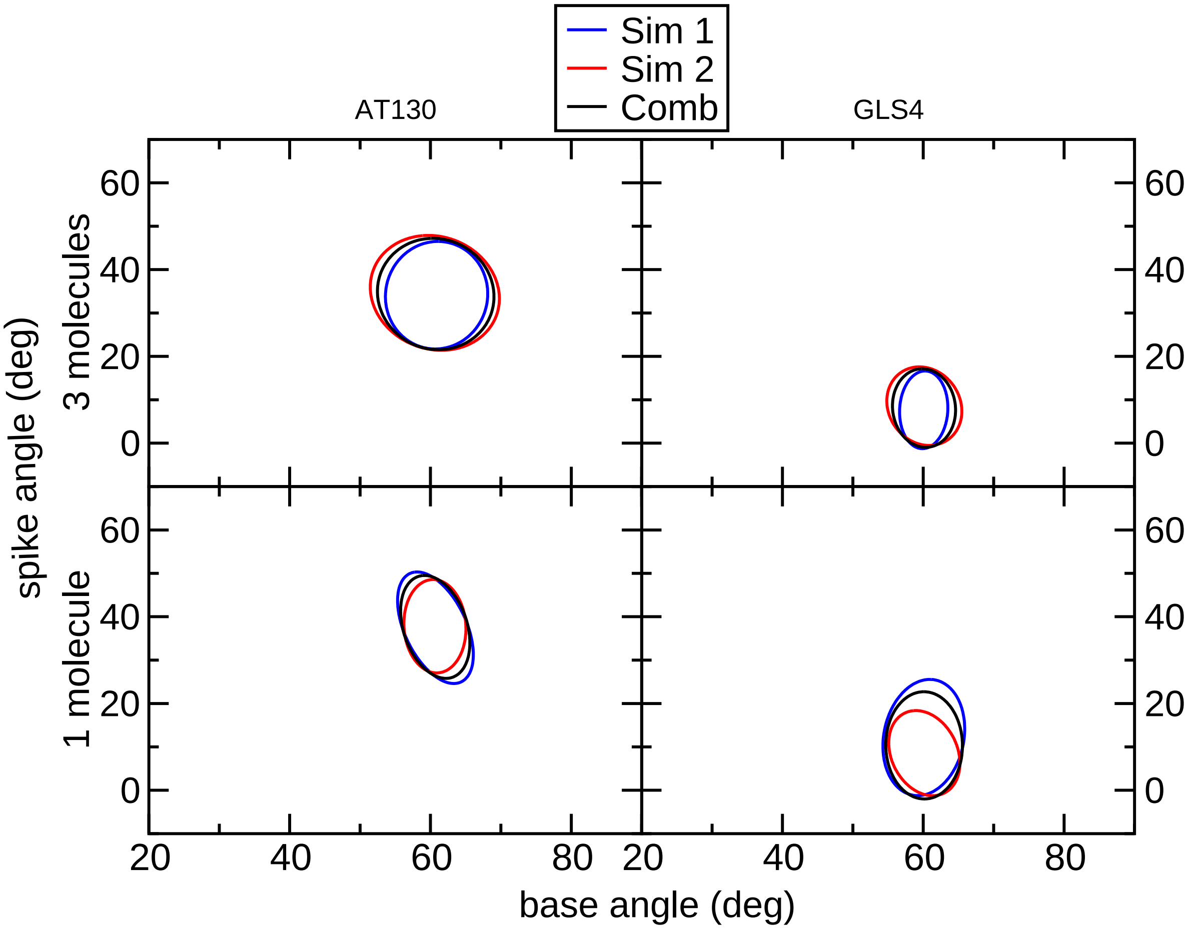


**Figure S23.** Comparison of standard deviation ellipses resulting from each of the two simulations to the one obtained from combining both simulations for hexamers with bound GLS4 and AT130. Top graphs display the results with 3 bound molecule. Bottom graphs show the results for hexamers with one bound molecule.

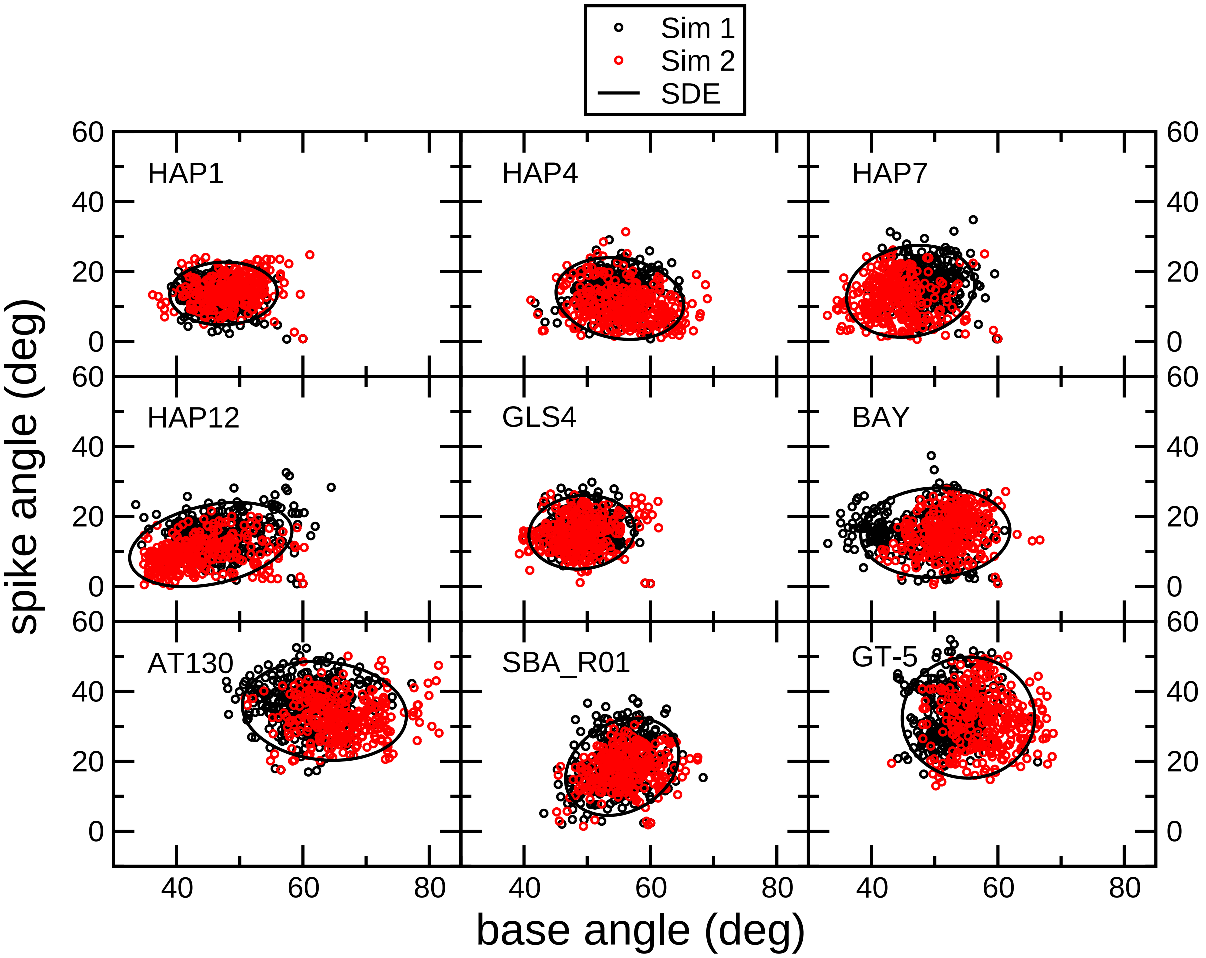


**Figure S24.** Distribution of spike and base angles and the resulting standard deviation ellipses for the studied tetramers with bound drug compounds.

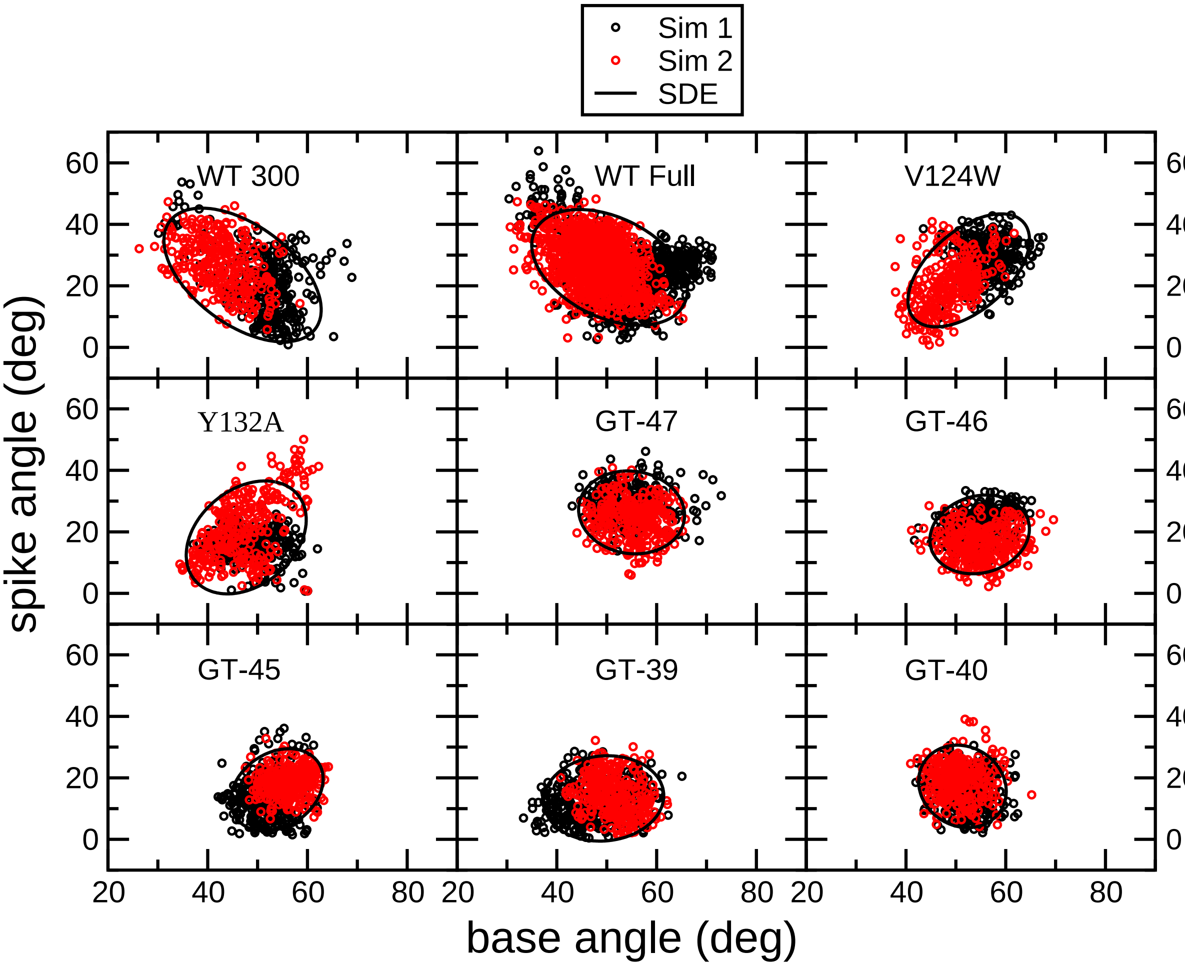


**Figure S25.** Distribution of spike and base angles and the resulting standard deviation ellipses for the studied apo-tetramers and tetramers with bound novel compounds.

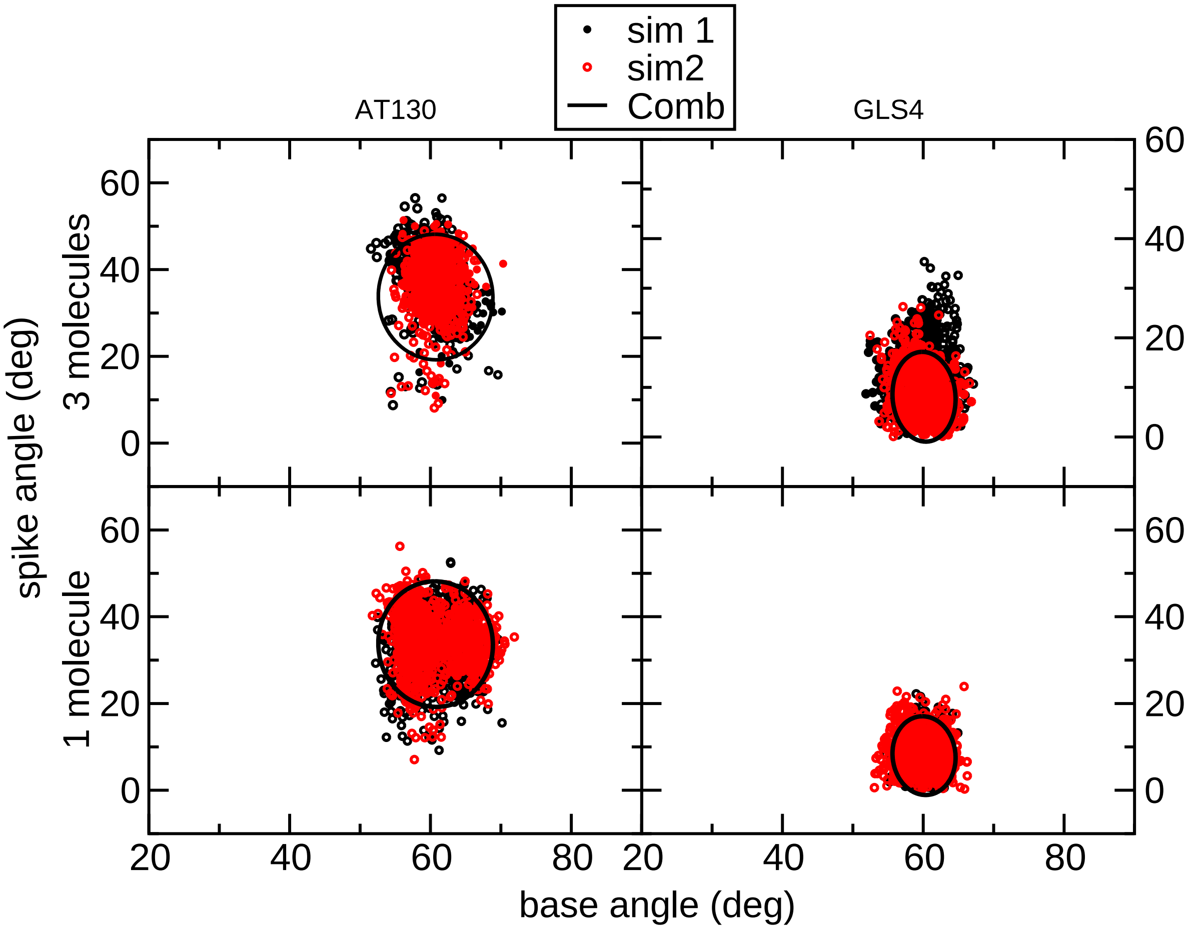


**Figure S26.** Distribution of spike and base angles and the resulting standard deviation ellipses for hexamers with bound GLS4 and AT130. Top graphs display the results with 3 bound molecule. Bottom graphs show the results for hexamers with one bound molecule.

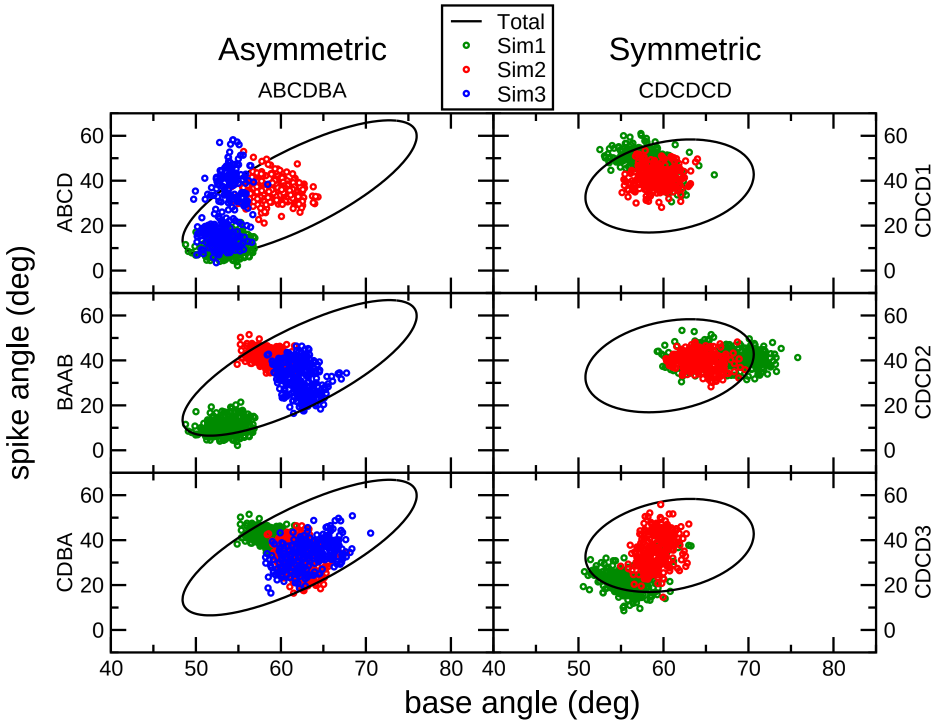


**Figure S27.** Distribution of spike and base angles for the different tetramers in the asymmetric (left graphs) and symmetric (right graphs) hexamer. Each graph shows the ellipses for a specific tetramer during different simulations and the ellipse resulting from averages of all tetramers and all simulations (black).

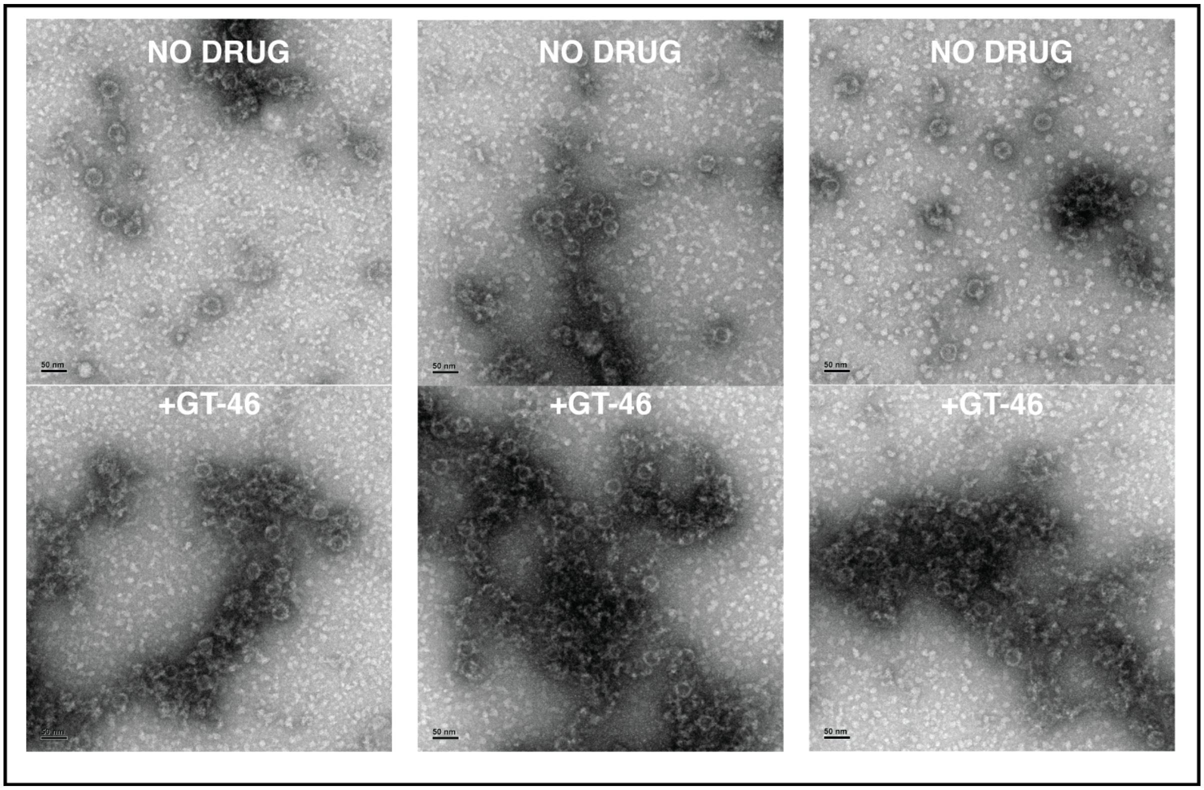


**Figure S28.** Full EM images with and without GT-46 present at 40 $\mu$M (top and bottom, respectively). See also Figure 5 in the main text.
